# Lineage-specific regulatory changes in the pathological cardiac remodeling of hypertrophy cardiomyopathy unraveled by single-nucleus RNA-seq and spatial transcriptomics

**DOI:** 10.1101/2021.07.21.453191

**Authors:** Xuanyu Liu, Kunlun Yin, Liang Chen, Wen Chen, Wenke Li, Taojun Zhang, Yang Sun, Meng Yuan, Hongyue Wang, Shuiyun Wang, Shengshou Hu, Zhou Zhou

**Affiliations:** State Key Laboratory of Cardiovascular Disease, National Center for Cardiovascular Diseases, Fuwai Hospital, the Chinese Academy of Medical Sciences, Beijing 100037, China; Center of Laboratory Medicine, Beijing Key Laboratory for Molecular Diagnostics of Cardiovascular Diseases, Fuwai Hospital, the Chinese Academy of Medical Sciences, Beijing 100037, China; Department of Cardiovascular Surgery, Fuwai Hospital, the Chinese Academy of Medical Sciences, Beijing 100037, China

**Author notes:** Dr. Liu, Dr. Yin and Dr. Chen contributed equally to this manuscript. Correspondence author: Dr. Zhou Zhou,; Dr. Shuiyun Wang; Dr. Shengshou Hu.

**Keywords:** hypertrophy cardiomyopathy, pathological remodeling, cardiac fibrosis, single-nucleus RNA-seq, spatial transcriptomics

## Abstract

**BACKGROUND:** Hypertrophy cardiomyopathy (HCM) is the most common cardiac genetic disorder with the histopathological features of cardiomyocyte hypertrophy and cardiac fibrosis. The pathological remodeling that occurs in the myocardium of HCM patients may ultimately progress to heart failure and death. A thorough understanding of the cell type-specific changes in the pathological cardiac remodeling of HCM is crucial for developing successful medical therapies to prevent or mitigate the progression of this disease.

**METHODS:** We performed single-nucleus RNA-seq of the cardiac tissues from 10 HCM patients and 2 healthy donors, and conducted spatial transcriptomic assays of 4 cardiac tissue sections from 3 HCM patients. Comparative analyses were performed to explore the lineage-specific changes in expression profile, subpopulation composition and intercellular communication in the cardiac tissues of HCM patients. Based on the results of independent analyses including pseudotime ordering, differential expression analysis, and differential regulatory network analysis, we prioritized candidate therapeutic targets for mitigating the progression to heart failure or attenuating the cardiac fibrosis in HCM. Using the spatial transcriptomic data, we examined the spatial activity patterns of the key candidate genes, pathways and subpopulations.

**RESULTS:** Unbiased clustering of 55,122 nuclei from HCM and healthy conditions revealed 9 cell lineages and 28 clusters. Significant expansion of vascular-related lineages and contraction of cardiomyocytes, fibroblasts and myeloid cells in HCM were observed. The transcriptomic dynamics during the transition towards the failing state of cardiomyocytes in HCM were uncovered. Candidate target genes for mitigating the progression to heart failure in HCM were obtained such as *FGF12*, *IL31RA*, *BDNF*, *S100A1*, *CRYAB* and *PROS1*. The transcriptomic dynamics underlying the fibroblast activation were also uncovered, and candidate targets for attenuating the cardiac fibrosis in HCM were obtained such as *RUNX1*, *MEOX1*, *AEBP1*, *LEF1* and *NRXN3*.

**CONCLUSIONS:** We provided a comprehensive analysis of the lineage-specific regulatory changes in HCM. Our analysis identified a vast array of candidate therapeutic target genes and pathways to prevent or attenuate the pathological remodeling of HCM. Our datasets constitute a valuable resource to examine the lineage-specific expression changes of HCM at single-nucleus and spatial resolution. We developed a web-based interface (http://snsthcm.fwgenetics.org/) for further exploration.

## INTRODUCTION

Hypertrophy cardiomyopathy (HCM) is the most common cardiac genetic disorder with an estimated minimal prevalence of 1 in 200.^1^ HCM is also the leading cause of sudden cardiac deaths (SCDs) in young people, accounting for 36% of SCDs in young athletes.^2^ HCM is characterized by an increase in left ventricular wall thickness in the absence of another cardiac or systemic disease.^3^ The key histopathological hallmarks of HCM include cardiomyocyte hypertrophy and disarray as well as cardiac fibrosis.^4^ Pathological cardiac remodeling occurs in the myocardium of HCM patients,^5^ manifesting as cardiomyocyte dysfunction, increased fibroblast activation (fibrosis), chronic inflammation and cell death. If left untreated, the pathological remodeling may ultimately lead to adverse events including heart failure, arrhythmias and death. In recent years, significant efforts have been made to design therapeutic agents for HCM, for example, MYK-461 for inhibition of cardiac myosin ATPase.^6^ A thorough understanding of the cellular and molecular changes in the pathological cardiac remodeling of HCM is crucial for developing successful medical therapies to prevent or mitigate the progression of this disease.

The transcriptomic alterations in the cardiac tissue of HCM have previously been examined at the tissue level via bulk RNA-seq.^7,8^ However, cell type-specific changes could not be obtained from bulk data. Single-cell or single-nucleus RNA-seq (snRNA-seq) could overcome this limitation and allows unbiased dissection of the cellular changes at an unprecedented resolution. Given the large size of adult human cardiomyocytes, snRNA-seq has been successfully applied to dissect the heterogeneity of the adult human heart under healthy^9^ and diseased conditions, for example, myocardial infarction.^10^ However, there is still a lack of research exploring the transcriptomic changes of the HCM in a single-nucleus resolution. The recent advent of spatially resolved transcriptomics has greatly expanded our scope and power to understand the cellular mechanism of diseases by providing spatial information of expression that is lost in single-cell/nucleus data.^11^ Integrated analysis of snRNA-seq and spatial transcriptomic data would profoundly improve our knowledge regarding the pathogenesis of diseases.

In this study, we performed snRNA-seq of the cardiac tissues from HCM patients and healthy donors. We also conducted spatial transcriptomic assays of cardiac tissue sections from HCM patients. Comparative analyses were performed to explore the lineage-specific changes in expression profile, subpopulation composition and intercellular communication in the cardiac tissues of HCM patients. We identified the transcriptomic dynamics during the transition towards the failing state of cardiomyocytes in HCM, and prioritized the candidate therapeutic target genes for mitigating the progression to heart failure in HCM, such as *FGF12*, *IL31RA*, *BDNF*, *S100A1*, *CRYAB* and *PROS1*. We also reconstructed the trajectory of fibroblast activation and prioritized the candidate targets for attenuating the cardiac fibrosis in HCM, such as *RUNX1*, *MEOX1*, *AEBP1*, *LEF1* and *NRXN3*. We provided a vast array of candidate target genes and pathways for designing therapeutic agents to prevent or attenuate the pathological remodeling of HCM. Our datasets constitute a valuable resource and we developed a web-based interface (http://snsthcm.fwgenetics.org/) for further exploration.

## METHODS

The data, analytic methods and materials will be made available on request only for the purposes of reproducing the results.

### Ethics statement

The recruitment of all subjects complied with the ethical regulations approved by the ethics committee of Fuwai Hospital, the Chinese Academy of Sciences (No. 2020-1315). Written informed consent was received from each patient.

### Study subject enrollment and cardiac tissue collection

The HCM patients (n=13) enrolled in this study underwent surgical myectomy in Fuwai Hospital from 2015 to 2021. All the patients met the diagnostic criteria^12^ for HCM with a maximal left ventricular wall thickness ≥ 15 mm or ≥ 13 mm in patients with a family history. All the patients belonged to the basal septum subtype, the most common and severe morphological subtype,^4^ in which cardiac hypertrophy mainly confines to the basal interventricular septum (IVS) adjacent to the aortic valve. All the patients exhibited left ventricular outflow tract (LVOT) obstruction (LVOT gradient ≥30 mm Hg at rest or on provocation). Patients were excluded if they had cardiac hypertrophy caused by secondary factors, including systemic hypertension, myocardial infarction, valvular disease or hemodynamic obstruction caused by left-sided obstructive lesions (e.g., valvular stenosis). In addition, patients were excluded if they had myocarditis and systemic disorders such as RASopathies, mitochondrial myopathies and storage diseases. For snRNA-seq, cardiac IVS tissues obtained from HCM patients (n=10) during surgical resection at the obstruction site were immediately frozen and stored in liquid nitrogen until use for nuclei isolation. For spatial transcriptomic assays, fresh cardiac IVS tissues from HCM patients (n=3) were concurrently frozen in isopentane precooled by liquid nitrogen and embedded in optical cutting tissue (OCT) compound. As a control for snRNA-seq, cardiac IVS tissues were obtained from healthy donors of heart transplants (n=2). Detailed methods are provided in the Extended Methods of the Data Supplement.

## RESULTS

### Single-nucleus and spatial transcriptomic sequencing of the cardiac IVS tissues from HCM patients and healthy donors

As illustrated in Figure 1A, the cardiac IVS tissues of HCM patients who underwent surgical myectomy were collected for snRNA-seq (n=10; 10 samples) and spatial transcriptomic assays (n=3; 4 tissue sections, of which HCM1220B and HCM1220C were from the same patient). As a control group (referred to as HEALTHY), cardiac IVS tissues from healthy donors of heart transplants (n=2; 3 samples, of which HEALTHY1A and HEALTHY1B were from the same donor) were also subjected to snRNA-seq. The control group was ethnicity-and sex-matched with the HCM group (Chinese, male). The detailed demographic and clinical information of the enrolled subjects were in Table I in the Data Supplement. After quality control, a total of 55,122 nuclei (HCM: 39,183; HEALTHY: 15,939) were obtained (Table II in the Data Supplement). For the spatial transcriptomic data, 3,339 to 4,849 spots were detected to be over tissue on the four sections (Table III in the Data Supplement). We developed a web-based interface (http://snsthcm.fwgenetics.org/) for all the datasets, which permit interactive examination of the expression of any gene or the activity of any pathway for both the snRNA-seq and spatial transcriptomic data.

**Figure 1.**
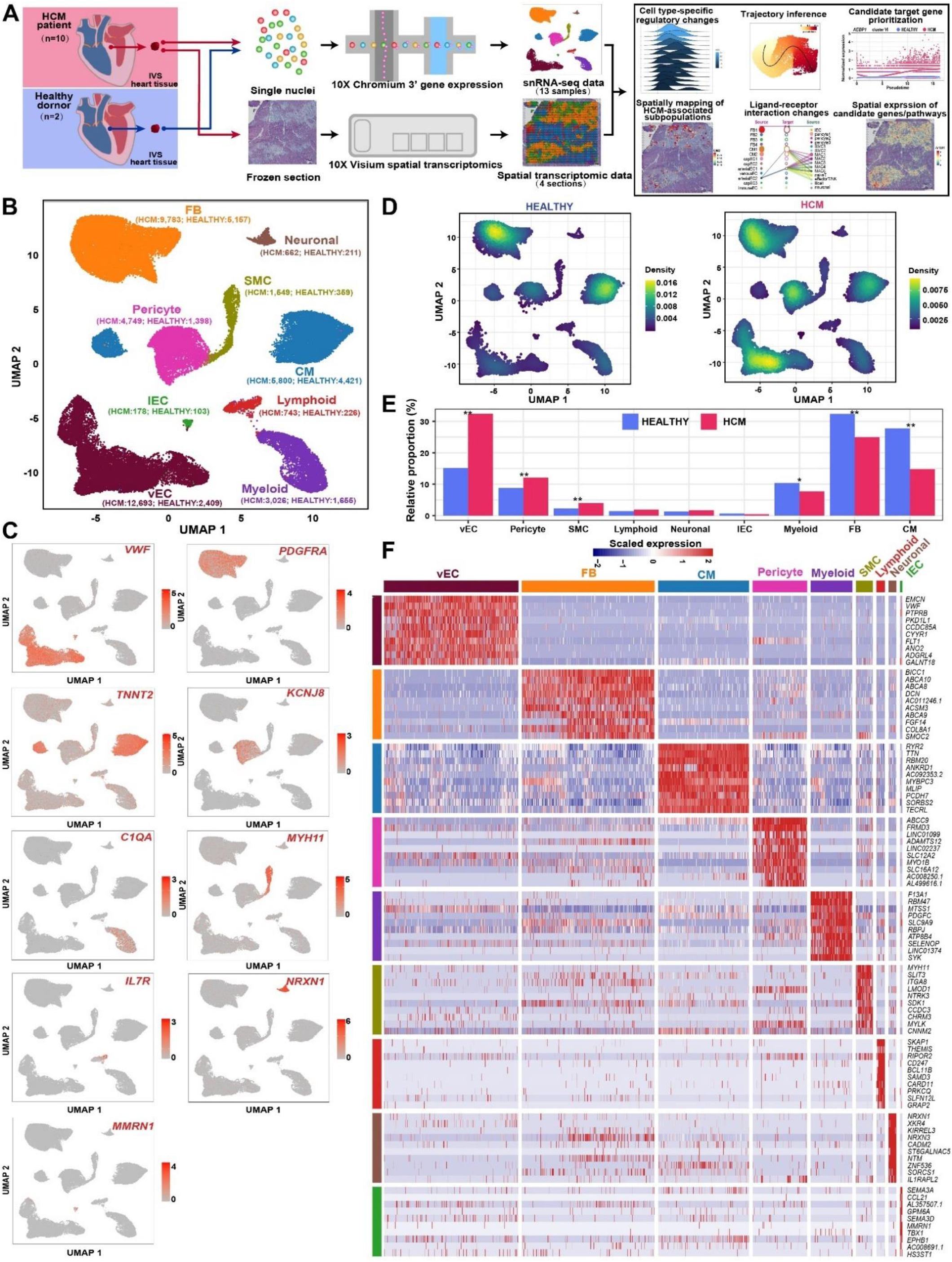
The proportional changes of the major cell types of the human cardiac tissues under HCM versus healthy conditions. **A,** Schematic representation of the overall experimental procedure. The cardiac IVS tissues of HCM patients who underwent surgical myectomy were collected for snRNA-seq (n=10;10 samples) and spatial transcriptomic assays (n=3; 4 tissue sections). As a control, cardiac IVS tissues from healthy donors of heart transplants (n=2; 3 samples) were subjected to snRNA-seq. **B,** Unbiased clustering of 55,122 nuclei from all 13 samples identifies 9 major cell types. The number in the parenthesis indicates the nucleus count. **C,** UMAP plots showing the expression of the established marker genes for each cell type. **D,** Comparison of the nucleus densities in the UMAP space between two conditions reveals remarkable changes in the relative proportion of cell types in HCM. Nuclei were randomly sampled for equal numbers in each group (n= 15,939). **E,** The relative proportion of each cell type in each condition. *: P-value < 0.05; **: P-value < 0.01. A permutation-based statistical test (differential proportion analysis; DPA). CM: cardiomyocyte; FB, fibroblast; lEC: lymphatic endothelial cell; SMC: smooth muscle cell; vEC: vascular endothelial cell.

### Significant expansion of vascular-related lineages and contraction of cardiomyocytes, fibroblasts and myeloid cells in HCM

Based on the expression of established markers for each lineage,^9,13^ as shown in Figure 1B and 1C, a total of 9 cell types were identified by joint clustering of the snRNA-seq data from both conditions: vascular endothelial cells (vECs, marked by *VWF*), fibroblasts (FBs, marked by *PDGFRA*), cardiomyocytes (CMs, marked by *TNNT2*), pericytes (marked by *KCNJ8*), myeloid cells (marked by *C1QA*), smooth muscle cells (SMCs, marked by *MYH11*), lymphoid cells (marked by IL7R), neuronal cells (marked by *NRXN1*) and lymphatic endothelial cells (lECs, marked by *MMRN1*). By comparing the nucleus densities in the UMAP space between the two conditions, remarkable changes in the relative proportion of cell types in HCM could be found, particularly for vECs, pericytes and cardiomyocytes (Figure 1D, Figure I in the Data Supplement). Next, we quantified the changes in cellular composition between the two conditions (Figure 1E). To determine whether the changes were expected by chance, we performed a permutation-based statistical test (differential proportion analysis; DPA) as described previously. ^14^ Vascular-related lineages including vECs, pericytes and SMCs were significantly expanded (P-value < 0.05, the DPA test), which was consistent with the knowledge of increased angiogenesis in HCM.^15^ Cardiomyocytes, fibroblasts and myeloid cells were significantly contracted (P-value < 0.05, the DPA test), which may reflect the increased cell death in HCM. Figure 1F shows the distinct molecular signatures of each lineage. To facilitate further data usage, the mean expression of all genes in each lineage under both conditions was provided in Table IV in the Data Supplement.

### Cardiomyocyte-specific regulatory changes in the pathological cardiac remodeling of HCM

Unbiased clustering grouped the cardiomyocytes into two subpopulations: CM1 and CM2 (Figure 2A; Table V in the Data Supplement). CM2 expressed high levels of maladaptive markers indicating the reactivation of the fetal gene program such as *NPPB* (encoding natriuretic peptide B, a clinically used biomarker for heart failure) and *ACTA1* (encoding skeletal α-actin),^16^ thus representing a failing state of cardiomyocytes (Figure 2B). CM1 expressed high levels of *FGF12* and *CORIN*, which may represent cardiomyocytes in a relatively homeostatic or compensatory hypertrophy state. Consistent with this, CM2 was significantly expanded in HCM, while CM1 was significantly contracted (Figure 2C; P-value < 0.01, the DPA test). Next, using DEsingle,^17^ we detected the differentially expressed genes in HCM versus HEALTHY in each lineage (Table VI in the Data Supplement). For cardiomyocytes, 2,021 genes were significantly upregulated, and 486 genes were significantly downregulated (the absolute of log2 fold change >1, adjusted P-value < 0.05). In agreement with the pathological hypertrophy phenotype of HCM, the upregulated genes were enriched for terms associated with cell growth and protein synthesis (e.g., “Ribosome assembly” and “Translation”), energy metabolism (e.g., “Oxidative phosphorylation”), stress response (e.g., “Cellular responses to stress”), immune response (e.g., “Antigen processing and presentation”), cell death (e.g., “Regulation of programmed cell death”), metabolic reprogramming (e.g., “Organonitrogen compound metabolic process”), as well as contraction (e.g., “Cardiac muscle contraction”; Figure 2D and Table VII in the Data Supplement). We further explored the dysregulated pathways in each lineage through gene set enrichment analysis (GSEA),^18^ which facilitates biological interpretation by robustly detecting concordant differences at the pathway level (Table VIII in the Data Supplement). As shown in Figure 2E, besides the pathways identified above by functional enrichment analysis, GASE analysis revealed more pathways that were upregulated in cardiomyocytes of HCM, for example, “NOTCH2 ACTIVATION AND TRANSMISSION OF SIGNAL TO THE NUCLEUS”, which supports the potential role of NOTCH signaling in cardiac hypertrophy. ^19^ In addition, using the method implemented in bigScale2,^20^ gene regulatory networks (GRNs) for each lineage were built separately for each condition. Comparative analysis of the GRNs between HCM and HEALTHY (differential regulatory networks analysis; DRN analysis) was performed for each lineage, and genes were ranked based on the changes in centrality, i.e., biological importance in the GRN (Table IX in the Data Supplement). Figure II in the Data Supplement shows the GRNs of cardiomyocytes in both conditions, and representative genes with great changes in centrality were identified and labeled such as *CRYAB* (Crystallin Alpha B), *EIF1* (Eukaryotic Translation Initiation Factor 1), *S100A1* (S100 Calcium Binding Protein A1), *PROS1* (Protein S), *TGFB2* (Transforming Growth Factor Beta 2) and *CREB5* (CAMP Responsive Element Binding Protein 5).

**Figure 2.**
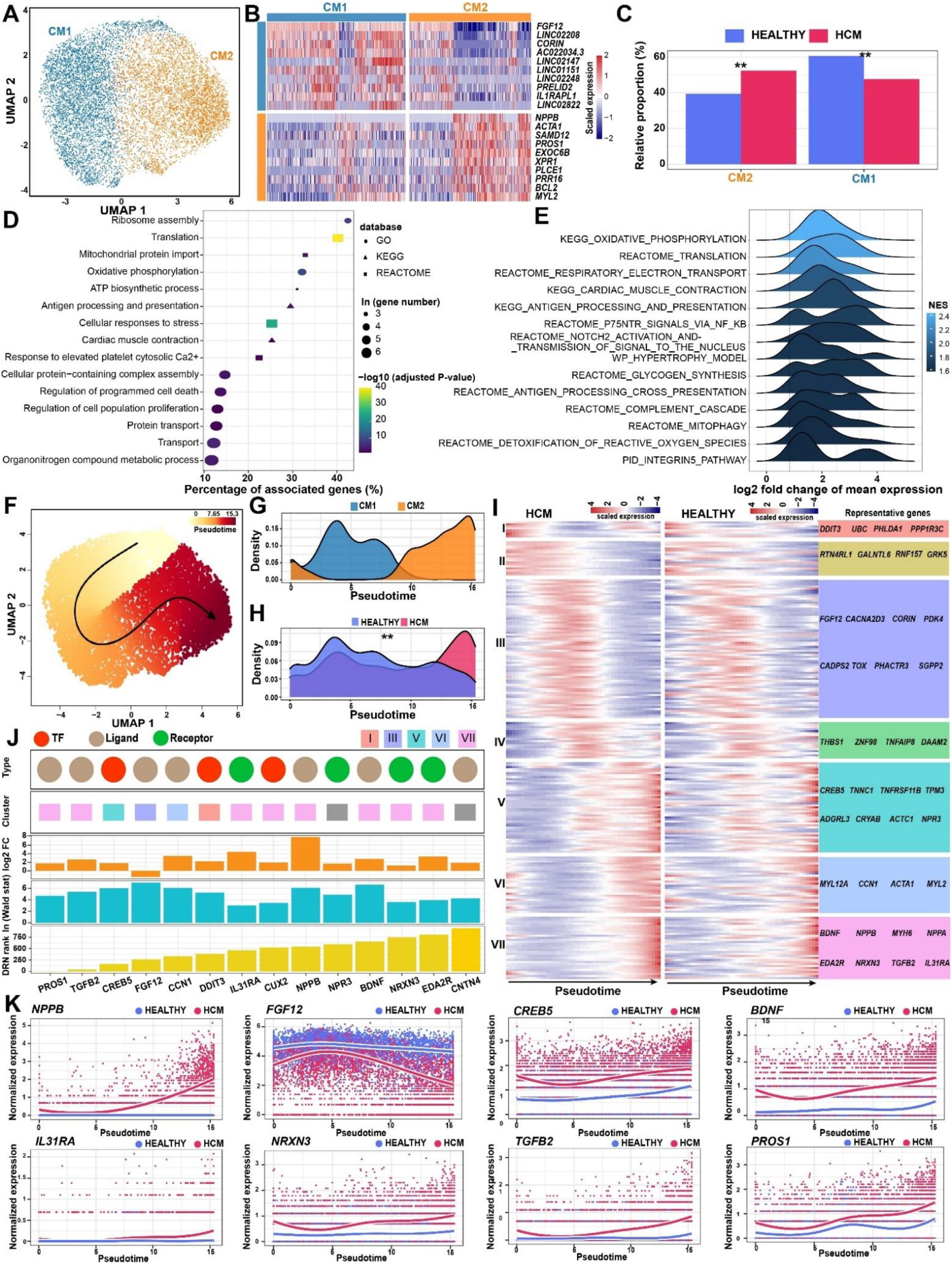
Cardiomyocyte-specific regulatory changes in the pathological remodeling of HCM. **A,** Subpopulations of cardiomyocytes. **B,** Heatmap showing the molecular signatures of each subpopulation. **C,** The relative proportion of each subpopulation in each condition. **: P-value of the permutation-based DPA test < 0.01. **D,** Representative terms enriched in the significantly upregulated genes in the cardiomyocytes of HCM. The significance threshold of the hypergeometric test was set to be an adjusted P-value < 0.05. **E,** Representative pathways that were significantly upregulated in the cardiomyocytes of HCM detected by GSEA. The density curve of the log2 fold change in the expression of the core enrichment genes for each pathway is shown. NES, normalized enrichment score. **F,** Cellular trajectory reconstructed for the transition towards failing cardiomyocytes in HCM using Slingshot. The arrow shows the direction of cellular state changes. **G,** Density curves showing the distributions of the two subpopulations along the trajectory. **H,** Density curves showing the distributions of the cardiomyocytes from different conditions along the trajectory. ** P-value < 2.2e-16; Kolmogorov-Smirnov test. **I,** Heatmaps showing the expression dynamics of the 216 genes with significantly different patterns along the trajectory between the two conditions. These genes were detected by differential expression pattern analysis using the “conditionTest” function of tradeSeq and were categorized into 7 gene clusters by hierarchical clustering. The significance threshold was set to be an adjusted P-value < 0.05. **J,** The candidate target genes that were prioritized based on the results of three independent analyses including the difference in expression patterns, the fold change of expression levels, and the centrality change in GRNs. DRN rank: the gene ranking based on the centrality change in GRNs obtained by differential regulatory network analysis. Log2FC: log2 fold change of the expression levels in cardiomyocytes. Wald stat: the statistics of differential expression pattern analysis. Only genes encoding transcription factors (TFs), ligands and receptors were considered. **K,** Smoothed expression curves of representative candidate targets along the trajectory in both conditions.

### Transcriptomic dynamics during the transition towards the failing state of cardiomyocytes in HCM

To decipher the transcriptomic dynamics during the transition towards the failing state of cardiomyocytes and identify potential targets for mitigating the progression of heart failure in HCM, we reconstructed the trajectory through the pseudo-temporal ordering of the nuclei of cardiomyocytes using Slingshot^21^ (Figure 2F). The failing cardiomyocytes of CM2 were ordered at relatively later pseudotime (Figure 2G). Significant differences existed between the pseudotime distributions of the two conditions (Figure 2H; P-value < 2.2e-16, the Kolmogorov-Smirnov test). Then, using tradeSeq,^22^ the genes with significantly different expression patterns along the trajectory between the two conditions were identified and clustered into 7 gene clusters (Figure 2I; Table X in the Data Supplement; adjusted P-value adjusted < 0.05). Notably, the maladaptive markers *NPPB* and *NPPA* were within the last gene cluster (VII). Next, we prioritized the candidate target genes for medical therapies based on the results of three independent analyses including the difference in expression patterns along the trajectory (adjusted P-value < 0.05), the fold change of expression levels between conditions (the absolute of log2 fold change > 1), and the centrality change in GRNs (DRN rank < 1000). Only genes encoding transcription factors (TFs), ligands and receptors were considered. Figure 2J showed 14 candidate genes we prioritized. For most of the genes, the roles in the transition of cardiomyocytes towards failing states in HCM have not been recognized previously such as *FGF12* (fibroblast growth factor 12), *CREB5*, *BDNF* (brain-derived neurotrophic factor), *IL31RA* (interleukin 31 receptor A), *NRXN3* (neurexin 3), *TGFB2* and *PROS1* (Figure 2K). Notably, some of them, e.g., *FGF12*, *IL31RA* and *PROS1* were significantly upregulated in the cardiac tissues of HCM (q-value < 0.05) according to the results of bulk RNA-seq^7^ previously performed by our lab (Figure III in the Data Supplement), further reflecting their roles in the pathogenesis of HCM.

### Fibroblast-specific regulatory changes in the pathological cardiac remodeling of HCM

Four fibroblast subpopulations were identified through unbiased clustering: *KCNMB2* ^high^ FB1, *NRXN3* ^high^ FB2, *CNTNAP2* ^high^ FB3 and *CD55* ^high^ FB4 (Figure 3A and 3B; Table V in the Data Supplement). Notably, FB2 expressed the highest levels of markers for activated fibroblasts (previously known as myofibroblasts^23^) such as *CCN2*, *FN1*, *COL1A1*, *COL3A1* and *MYH10,*^24^ thus representing an activated state of fibroblasts. Hierarchical clustering revealed a close relationship between FB1 and FB2 (Figure 3D), and thus FB1 may represent a state of quiescent fibroblasts. Consistent with this, FB2 was significantly expanded while FB1 was significantly contracted in HCM versus HEALTHY (Figure 3E; P-value < 0.01, the DPA test). Next, differentially expressed genes in fibroblasts between the two conditions were detected (Table VI in the Data Supplement). In line with the fibrosis that occurred in HCM, fibrosis-associated terms such as “Extracellular matrix organization” and “Cellular response to transforming growth factor beta stimulus” were enriched in the upregulated genes (Figure 3F and Table VII in the Data Supplement). In addition, the upregulated genes were also enriched for terms related to protein translation and processing, energy metabolism, stress response, as well as immune response. Notably, Hedgehog signaling and G protein-coupled receptor (GPCR) signaling were also enriched (Figure 3F), consistent with their roles in fibrogenesis known in other tissues and disease conditions.^25,26^ Moreover, GSEA revealed a more comprehensive list of signaling pathways that were upregulated in fibroblasts of HCM (Figure 3G and Table VIII in the Data Supplement), including classic profibrotic signaling pathways^27^ (e.g., “TGF BETA SIGNALING PATHWAY” and “WNT SIGNALING PATHWAY”) and cell surface ECM receptor pathways (e.g., “SYNDECAN 1 PATHWAY” and “INTEGRIN A4B1 PATHWAY”). In addition, the top five genes with great changes in centrality that were detected through DRN analysis included *ADAM19* (ADAM metallopeptidase domain 19), *RUNX1* (RUNX family transcription factor 1), *CTIF* (cap-binding complex dependent translation initiation factor), *MEOX1* (mesenchyme homeobox 1) and *FGF7* (fibroblast growth factor 7; Figure IV and Table IX in the Data Supplement).

**Figure 3.**
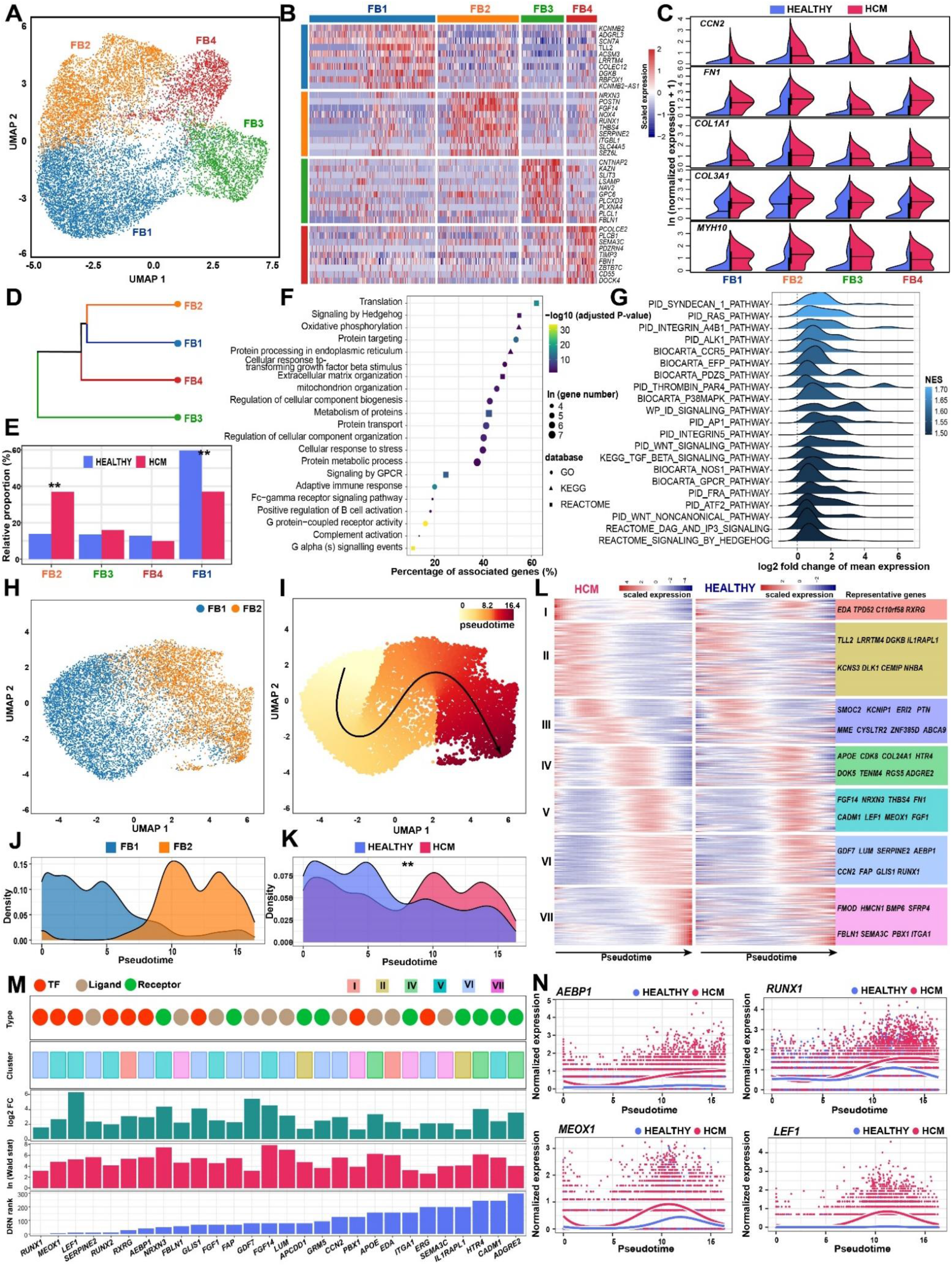
Fibroblast-specific regulatory changes in the pathological remodeling of HCM. **A,** Subpopulations of fibroblasts. **B,** Heatmap showing the molecular signature of each subpopulation. **C,** Split violin plots showing the expression of the markers for activated fibroblasts. **D,** Hierarchical clustering of the subpopulations. **E,** The relative proportion of each subpopulation in each condition. **: P-value of the permutation-based DPA test < 0.01. **F,** Representative terms enriched in the significantly upregulated genes in the fibroblasts of HCM. The significance threshold of the hypergeometric test was set to be an adjusted P-value < 0.05. **G,** Representative pathways that were significantly upregulated in the fibroblasts of HCM detected by GSEA. The density curve of the log2 fold change in the expression of the core enrichment genes for each pathway is shown. NES, normalized enrichment score. **H,** UMAP plot showing the subpopulations FB1 and FB2. **I,** Cellular trajectory reconstructed for the fibroblast activation in HCM using Slingshot. The arrow shows the direction of cellular state changes. **J,** Density curves showing the distributions of the two subpopulations along the trajectory. **K,** Density curves showing the distributions of the fibroblasts from different conditions along the trajectory. ** P-value < 2.2e-16; Kolmogorov-Smirnov test. **L,** Heatmaps showing the expression dynamics of the 432 genes with significantly different patterns along the trajectory between the two conditions. These genes were detected by differential expression pattern analysis using the “conditionTest” function of tradeSeq and were categorized into 7 gene clusters by hierarchical clustering. The significance threshold was set to be an adjusted P-value < 0.05. **M,** The candidate target genes that were prioritized based on the results of three independent analyses including the difference in expression patterns, the fold change of expression levels, and the centrality change in GRNs. **N,** Smoothed expression curves of representative candidate targets along the trajectory in both conditions.

### Transcriptomic dynamics during the activation of fibroblasts in HCM

To decipher the transcriptomic dynamics during the activation of fibroblasts and identify candidate therapeutic targets to alleviate the cardiac fibrosis in HCM, we reconstructed the trajectory of fibroblast activation through the pseudo-temporal ordering of the nuclei of FB1 and FB2 (Figure 3H and 3I). The activated fibroblasts FB2 were ordered at the end of the trajectory (Figure 3J). The pseudotime distribution of the fibroblasts of HCM was significantly different from those of HEALTHY (Figure 3K; P-value < 2.2e-16, Kolmogorov-Smirnov test). Next, the genes exhibiting significantly different expression patterns along the trajectory between the two conditions were identified and clustered into 7 gene clusters (Figure 3L; Table X in the Data Supplement; P-value adjusted for multiple testing < 0.05). Then, we prioritized the candidate target genes according to the criteria described above. Figure 3M showed 28 candidate genes that we prioritized. Notably, the top candidates included TF genes such as *RUNX1*, *MEOX1*, *LEF1* (lymphoid enhancer-binding factor 1) and *AEBP1* (AE Binding Protein 1), which were significantly more upregulated along the trajectory of fibroblast activation in HCM versus HEALTHY (Figure 3N). In addition, results from the bulk RNA-seq^7^ showed that some of the genes, such as *AEBP1*, *LEF1*, *NRXN3* and *GLIS1*, were significantly upregulated in the cardiac tissues of HCM, further reflecting their roles in the pathogenesis of HCM (Figure III in the Data Supplement).

### The subpopulations of the immune and vascular lineages and their proportional changes in HCM

Unbiased clustering revealed 8 immune subpopulations (Figure 4A). The subpopulations immune_c0, c1, c4, c5 and c6 expressed high levels of *CD68* (Figure 4B), thus representing five subpopulations of macrophages (which were referred to as MAC1-5 hereafter). As shown in Figure 4C, *FGF13* ^high^ MAC1 and *IGSF21* ^high^ MAC2 expressed high levels of *LYVE1*, which marked for vessel-associated resident macrophages with M2-like phenotypes.^28^ MAC5 expressed high levels of *FCN1,* which marks proinflammatory macrophages.^29^ Differential proportional analysis revealed a significant expansion of MAC2 and contraction of MAC1 (Figure 4D; P-value < 0.05, the DPA test), and thus MAC2 represented a more activated state compared to MAC1. Both functional enrichment analysis and GSEA supported the immune activation of macrophages in HCM (Figure VA in the Data Supplement). Besides a small cluster of the nuclei of B cells (marked by *CD79A*), another two closely related subpopulations of the lymphoid lineages were identified: immune_c2 and immune_c3. Immune_c2 expressed high levels of the T cell marker *CD3D* (Figure 4B) and exhibited high naiveness scores (Figure 4E), thus representing nuclei of naïve T cells. Immune_c3 expressed high levels of the T cell marker *CD3D* and the Natural Killer (NK) cell marker *NCR1*, and exhibited high cytotoxicity scores (Figure 4E), thus representing a mixture of the nuclei of effector T/NK cells. Expectedly, we observed a significant expansion of the effector T/NK nuclei and a significant contraction of the naïve T nuclei (Figure 4F; P-value < 0.05, the DPA test).

**Figure 4.**
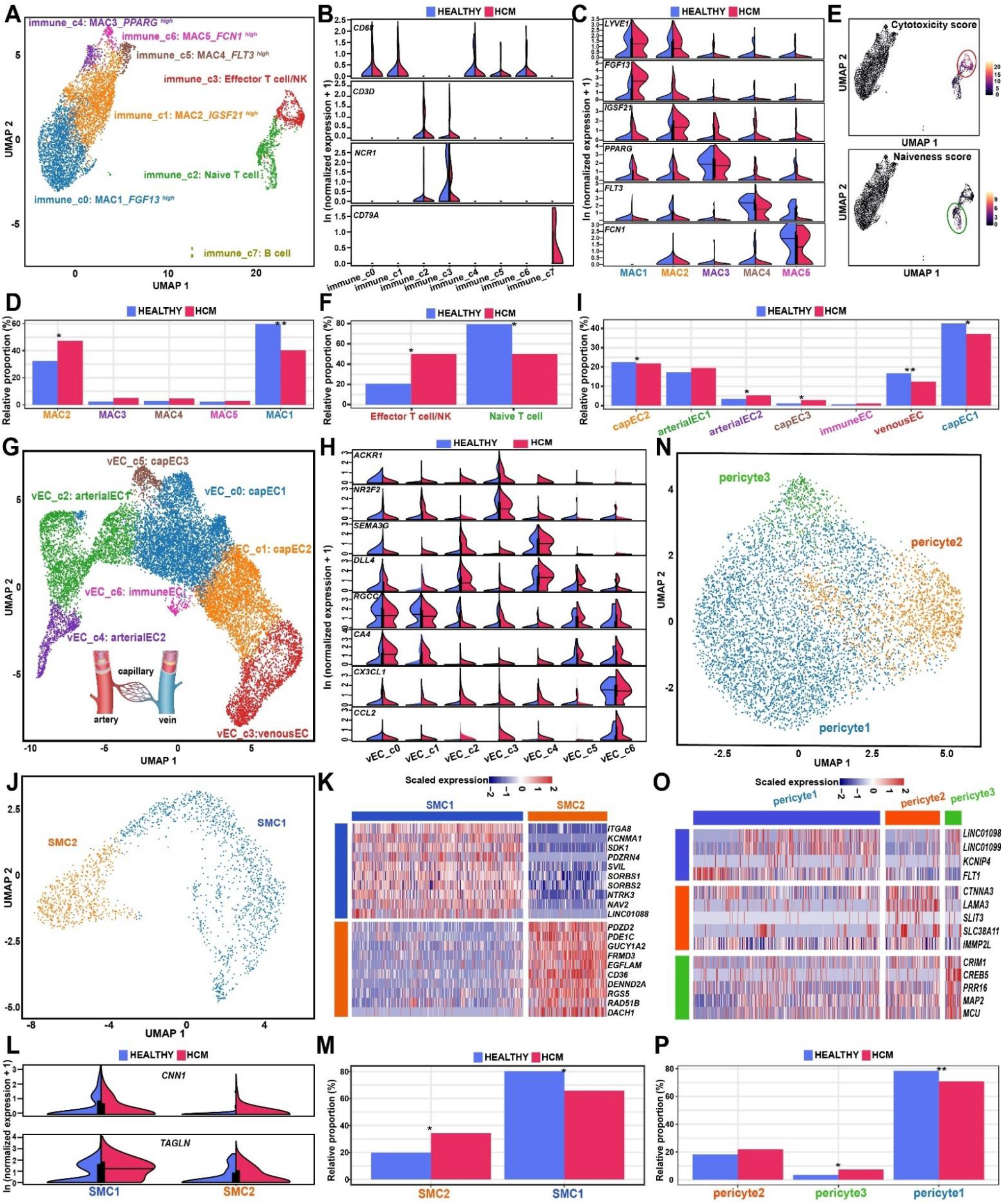
The subpopulations of the immune and vascular lineages and their proportional changes in HCM. **A**, UMAP plots showing the subpopulations of the immune lineage. B, Expression of the established markers for macrophages (*CD68*), T cells (*CD3D*), Natural killer cells (*NCR1*) and B cells (*CD79A*) in each immune subpopulation. **C,** Expression of the marker for each of the five macrophage subpopulations. **D,** Relative proportion of each subpopulation of macrophages in each condition. **E,** UMAP plots showing the cytotoxicity and naiveness scores for each immune nucleus. The cytotoxicity and naiveness scores were calculated by summing the expression of previously reported signatures for T cell cytotoxicity (*PRF1*, *IFNG*, *GNLY*, *NKG7*, *GZMB*, *GZMA*, *GZMH*, *KLRK1*, *KLRB1*, *KLRD1*, *CTSW* and *CST7*) and naiveness (*TCF7*, *SELL*, *LEF1* and *CCR7*).^48^ **F,** Relative proportion of each subpopulation of T/NK cells in each condition. **G,** UMAP plots showing the subpopulations of the vECs. **H,** Expression of the established markers for venous ECs (*ACKR1* and *NR2F2*), arterial ECs (*SEMA3G* and *DLL4*), capillary ECs (*RGCC* and *CA4*) and immune ECs (*CX3CL1* and *CCL2*). **I,** Relative proportion of each subcluster of the vECs in each condition. **J,** UMAP plots showing the subpopulations of the SMCs. **K,** Molecular signature for each SMC subpopulation. **L,** Expression of contractile markers *CNN1* and *TAGLN* in each SMC subpopulation. **M,** Relative proportion of each SMC subpopulation in each condition. **N,** UMAP plots showing the subpopulations of the pericytes. **O,** Molecular signature for each pericyte subpopulation. **P,** Relative proportion of each subpopulation of the pericytes in each condition. In E, F, I, M and P, *: P-value < 0.05, **: P-value < 0.01, the permutation-based DPA test. MAC: macrophage; SMC: smooth muscle cell; vEC: vascular endothelial cell.

For the vEC lineage, we identified 7 subpopulations that were aligned consecutively in the UMAP space (Figure 4G). From the left to the right of UMAP1, based on the established markers,^9^ the subclusters were assigned to arterial ECs (marked by *SEMA3G* and *DLL4*; arterial EC2 and arterial EC1), capillary ECs (marked by *RGCC* and *CA4*; capEC3, capEC1, immune EC and capEC2) and venous ECs (marked by *ACKR1* and *NR2F2*; venousEC; Figure 4H). A significant expansion of most subpopulations except for capEC1 and venousEC was observed (P-value < 0.05, the DPA test; Figure 4I). For SMCs, two subpopulations were identified with distinct expression profiles: SMC1 and SMC2 (Figure 4J and 4K). Compared with SMC1, SMC2 expressed lower levels of contractile markers such as *CNN1* and *TAGLN* (Figure 4L) and were aligned closely to pericytes in the UMAP space (Figure VI in the Data Supplement). These results suggest that SMC2 may represent the vascular SMCs of the small vasculature that was greatly expanded in HCM. Consistent with this, a significant expansion of SMC2 was observed (P-value < 0.05, the DPA test; Figure 4M). For pericytes, three subpopulations were identified: pericyte1, pericyte2 and pericyte3 (Figure 4N and 4O), and an expansion of pericyte2 was observed (P-value < 0.05, the DPA test; Figure 4P). The subpopulation pericyte2 was closely related to SMCs based on the alignment in the UMAP space (Figure VI in the Data Supplement), which may represent pericytes surrounding the capillaries with relatively large caliber. The representative pathways upregulated in each of the three types of vascular lineage were shown in Figure V in the Data Supplement. Notably, like those observed in cardiomyocytes and fibroblasts, energy metabolism and immune response-related pathways were upregulated in all three cell types.

### Intercellular communication changes in the cardiac tissue of HCM inferred from the snRNA-seq data

To date, Intercellular interactions in HCM have mostly been characterized *in vitro* through coculture experiments. Based on the snRNA-seq data, CellChat ^30^ was used to infer ligand-receptor interactions among subpopulations *in vivo* separately for each condition (Table XI in the Data Supplement). Through pattern recognition approaches, the dominant incoming and outgoing signal patterns for each subpopulation were detected for each condition (Figure VII in the Data Supplement), and subpopulations belonging to the same lineage had more similar patterns. The inferred total number (Figure 5A) and strength of interactions (Figure 5B) were significantly increased in HCM, reflecting an enhanced intercellular communication in diseased conditions as reported in other diseases.^31^ Fibroblast subpopulations had a great increase in the number (Figure 5C) and strength (Figure 5D) of interactions for both outgoing and incoming signals, reflecting their central roles in the pathological remodeling of HCM. Notably, neuronal cells exhibited significantly enhanced incoming signals from other lineages, e.g., fibroblasts. Remarkably, cardiomyocytes, especially the failing subpopulation CM2, exhibited reduced communication with themselves (autocrine) and some other lineages (paracrine), e.g., macrophages. The remarkable change of cardiomyocytes in communication could also be observed by comparing the relative positions of cardiomyocytes in the 2D signal space between HEALTHY (Figure 5E) and HCM (Figure 5F).

**Figure 5.**
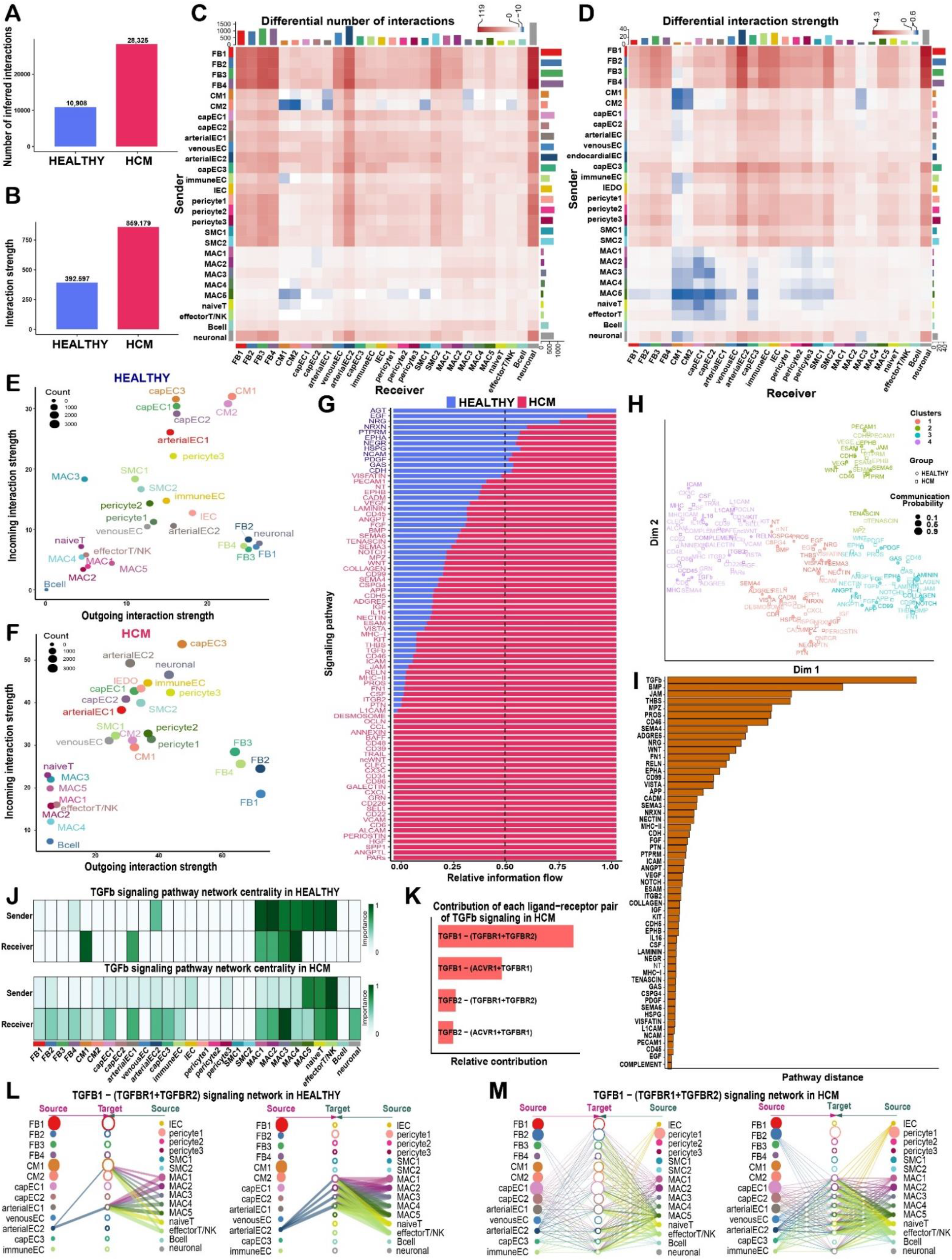
Intercellular communication changes in the cardiac tissues of HCM. **A,** Bar plot showing the total number of ligand-receptor interactions among the subpopulations of the cardiac tissues in both conditions. **B,** Bar plot showing the total interaction strength among the subpopulations of the cardiac tissues in both conditions. The total interaction strength was calculated by summing the communication probability of all inferred interactions. **C,** Heatmap showing the differential number of interactions among subpopulations in HCM versus HEALTHY. In the color bar, red represents an increase in the number of interactions and blue represents a decrease in the number of interactions. The top bar plot shows the sum of the changes in the number of incoming signals for each subpopulation. The right bar plot shows the sum of the changes in the number of outgoing signals for each subpopulation. **D,** Heatmap showing the differential interaction strength among subpopulations in HCM versus HEALTHY. **E,** Bubble plot showing the incoming and outgoing interaction strength for each subpopulation in HCM. The dot size represents the count of interactions. **F,** Bubble plot showing the incoming and outgoing interaction strength for each subpopulation in HEALTHY. **G,** Relative information flow for each signaling pathway in both conditions. The information flow is defined by the sum of the communication probability among all pairs of subpopulations. **H,** Joint manifold learning of the HCM and HEALTHY communication networks and grouping the signaling pathways based on functional similarity. A high degree of functional similarity means that the major senders and receivers are similar. **I,** The Euclidean distance of each pathway in the learn joint manifold. A larger distance means a larger difference in functional similarity (i.e., similarity in senders and receivers) between the two conditions. Only overlapping pathways between the two conditions are shown. **J,** The major senders and receivers of the TGFb signaling pathway inferred through network centrality analysis in HEALTHY (upper panel) and HCM (lower panel). **K,** Relative contribution of each ligand-receptor pair to the overall signal of the TGFb pathway in HCM. **L,** Hierarchical plot showing the inferred communication network for TGFB1-(TGFBR1+TGFBR2) signaling in HEALTHY. **M,** Hierarchical plot showing the inferred communication network for TGFB1-(TGFBR1+TGFBR2) signaling in HCM. In L and M, open and solid circles represent target and source, respectively. Edge width represents the interaction strength and circle size is proportional to the number of nuclei in each subpopulation. Edges are color-coded by the signal source.

Next, we compared the relative information flow for each signaling pathway between two conditions (Figure 5G; Figure VIII in the Data Supplement), and identified pathways that were greatly enhanced in HCM (e.g., PTN, ITGB2, CSF, PROS, ICAM, CD46, TGFb, MHC-1, ESAM and WNT) or specific to the HCM condition (e.g., PARs, ANGPTL and SPP1). Through joint manifold learning of the inferred communication networks, the signaling pathways were grouped based on functional similarity (i.e., similarity in senders and receivers, Figure 5H). The Euclidean distance of each pathway in the learn joint manifold reflected the changes in functional similarly between the two conditions. As shown in Figure 5I, the pathway with the largest distance was the TGFb pathway. In line with this, network centrality analysis confirmed that the TGFb pathway greatly changed in senders and receivers in HCM (Figure 5J): the top sender changed from MAC2 in HEALTHY to effector T/NK cells in HCM, and the top receiver changed from CM1 to MAC3. Then, we found that TGFB1-(TGFBR1+TGFBR2) was the ligand-receptor pair that contributed most to the network of TGFb signaling in the cardiac tissues of HCM (Figure 5K). As shown in Figure 5L and 5M, TGFB1-(TGFBR1+TGFBR2) signaling was enhanced in HCM, and the paracrine signal of TGFB1 received by fibroblasts, cardiomyocytes and vECs were predominately secreted by effector T/NK cells, naïve T cells and proinflammatory macrophages MAC5. The communication strength among subpopulations for any pathway or ligand-receptor pair is accessible through our web-based interface (http://snsthcm.fwgenetics.org/).

### Spatially resolved examination of the expression of candidate genes, the activity of HCM-associated pathways and subpopulations by spatial transcriptomics

As shown in Figure IX in the Data Supplement, the four tissue sections selected for spatial transcriptomic assays contained regions with replacement fibrosis and/or diffuse (interstitial or perivascular) fibrosis that commonly occur in HCM. The section HCM1225D was characterized by large replacement fibrotic scars and interstitial fibrosis (also see Figure 6A and 6B). For the section HCM1221A, interstitial fibrosis was restricted to a relatively narrow region close to the endocardium, while most regions represented non-fibrotic cardiac tissues. The sections HCM1220B and HCM1220C were featured by extensive diffuse fibrosis. Through unbiased clustering of the spots, spots in fibrotic regions could be separated from those in non-fibrotic regions. For example, spot clusters SC0 and SC1 generally represented spots in fibrotic and non-fibrotic regions on the section HCM1225D, respectively (Figure 6C and 6D; Figure X in the Data Supplement). Likewise, the fibrotic and non-fibrotic spot clusters were identified for other sections (Figure XI-XIII in the Data Supplement). Following the label transfer workflow of Seurat, we integrated the snRNA-seq data and the spatial transcriptomic data. CM1, the cardiomyocyte subpopulation in a homeostatic or compensatory hypertrophy state (marked by *FGF12*), was predicted to be localized in non-fibrotic regions, while CM2, the cardiomyocyte subpopulation in a failing state (marked by *NPPB*), was localized close to the fibrotic regions (Figure 6E and 6F). FB1, the quiescent fibroblast subpopulation, was mainly in non-fibrotic regions, while FB2, the activated fibroblast subpopulation, was in fibrotic regions (Figure 6E). The candidate target genes *AEBP1*, *RUNX1*, *MEOX1* and *MGP* were highly expressed in fibrotic regions (Figure 6F). Therefore, the spatial transcriptomic data confirmed the results of our snRNA-seq analysis above. Next, the dysregulated genes and pathways in fibrotic versus non-fibrotic regions were identified separately for each section (Table XII-XIII in the Data Supplement). As shown in Figure 6G, the upregulated pathways were mainly involved in ECM remodeling (e.g., “REACTOME_EXTRACELLULAR_MATRIX_ORGANIZATION” and “REACTOME_TRANSLATION”), fibrosis-related signaling (e.g., “KEGG_TGF_BETA_SIGNALING_PATHWAY” and “WP_PI3KAKT_SIGNALING_PATHWAY”) and immune response (e.g., “REACTOME_INTERFERON_SIGNALING” and “WP_IL18_SIGNALING PATHWAY”), while the downregulated pathways were mainly involved in contraction (e.g., “KEGG_CARDIAC_MUSCLE_CONTRACTION”), energy metabolism (e.g., “KEGG_OXIDATIVE PHOSPHORYLATION”) and TP53-mediated stress response (e.g., “REACTOME_TRANSCRIPTIONAL_REGULATION_BY_TP53”). Figure 6H showed that the representative upregulated pathways exhibited high expression activity in the fibrotic regions.

**Figure 6.**
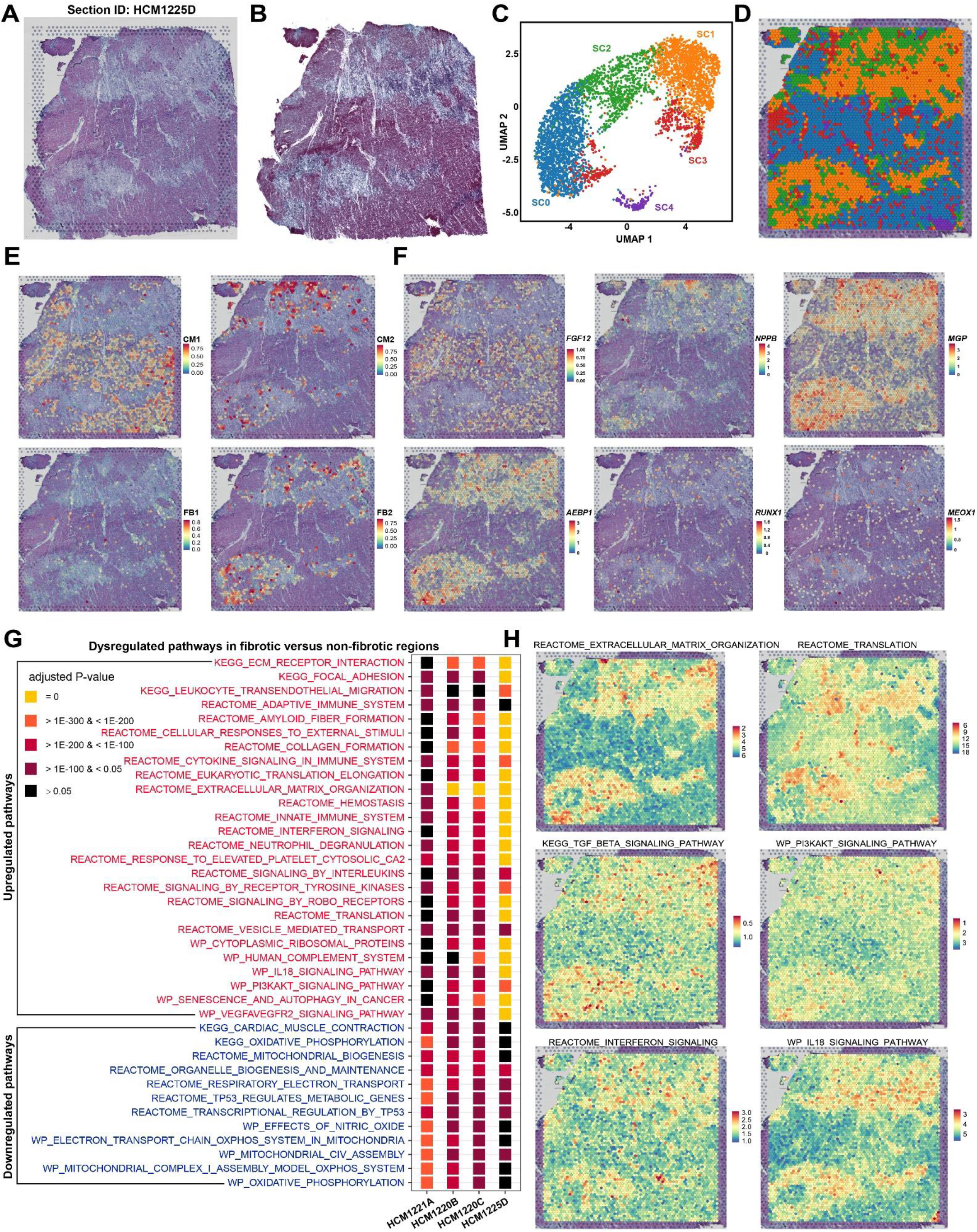
Spatially resolved examination of the expression of candidate genes, the activity of HCM-associated pathways and subpopulations by spatial transcriptomics. **A**, H&E staining image for the cardiac tissue section HCM1225D. **B**, Masson’s trichrome staining image for a section adjacent to HCM1225D. **C,** UMAP plot showing the spot clusters identified by unbiased clustering of the spots on HCM1225D. **D,** Distribution of the spot clusters on the section HCM1225D. **E,** Spatial location of the subpopulations FB1, FB2, CM1 and CM2 on the section HCM1225D predicted by integrating the snRNA-seq data and the spatial transcriptomic data. **F,** Expression distribution of representative markers and candidate target genes on the section HCM1225D. **G,** Dysregulated pathways in fibrotic versus non-fibrotic regions of the cardiac tissue sections of HCM. The dysregulated pathways were detected based on the pathway activity score of each spot using the Wilcoxon rank-sum test. The significance threshold was set to be a P-value adjusted for multiple testing < 0.05. The tests were performed separately for each of the four sections. **H,** Activity of representative upregulated pathways in fibrotic regions on the section HCM1225D.

## DISCUSSION

Understanding the regulatory changes under diseased conditions is of fundamental importance for successful drug development. By using snRNA-seq and spatial transcriptomic assays, the present study provided the first comprehensive analysis of the lineage-specific changes in expression profile, subpopulation composition and intercellular communication in the cardiac tissues of human HCM patients. The candidate genes we prioritized based on multiple independent analyses may serve as therapeutic targets to prevent or attenuate the pathological remodeling of HCM.

While cardiac remodeling is orchestrated by multiple lineages, the cardiomyocytes function as the most important determinants of cardiac conditions.^32^ Thus, medical therapies directly targeting the cardiomyocytes may represent the most promising strategy to alleviate pathological hypertrophy or mitigate the progression to heart failure in HCM. The single-nuclei resolution data allowed us to examine the cardiomyocyte-specific regulatory changes of HCM *in vivo* (Table VI in the Data Supplement). The results of the functional enrichment analysis and GSEA (Figure 2D and 2E) well recapitulated the features known for pathological cardiac hypertrophy,^16^ including increased protein translation, energy metabolism, stress response, immune response, cell death and contraction. Among the genes that were greatly changed in centrality through the DRN analysis (Figure II and Table IX in the Data Supplement), some have been implicated in cardiac hypertrophy or heart failure; For example, *CRYAB* (Crystallin Alpha B) has been demonstrated to suppress pressure overload-induced cardiac hypertrophy in mice.^33^ *S100A1* (S100 Calcium Binding Protein A1) has been suggested to be a target for the treatment of heart failure.^34^ However, the precise roles of most genes have not been elucidated in the pathogenesis of HCM such as *PROS1* (Protein S) and *CREB5* (CAMP Responsive Element Binding Protein 5). Cardiomyocytes were clustered into two subpopulations: CM1 and CM2 (Figure 2A), which represented a homeostatic or compensatory hypertrophy state and a failing state, respectively. The failing cardiomyocyte subpopulation CM2 was found to be close to the fibrotic regions by spatial transcriptomics (Figure 6E), which reflected the detrimental effects of cardiac fibrosis on cardiomyocytes in HCM. Intercellular communication analysis revealed that cardiomyocytes, especially the failing subpopulation CM2, exhibited reduced communication with themselves (autocrine) and some other lineages (paracrine) in HCM (Figure 5C and 5D), reflecting communication dysfunction of cardiomyocytes in HCM.

Pathological cardiac hypertrophy is a common predecessor to heart failure.^35^ A recent study reported the transcriptomic differences of cardiomyocytes between early (hypertrophic cardiomyocytes) and maladaptive phage (failing cardiomyocytes) of cardiac remodeling in pressure overload-induced mouse models,^36^ Through pseudo-temporal ordering, we identified the transcriptomic dynamics during the transition towards the failing state of cardiomyocytes in HCM of human patients (Figure 2I). Based on multiple lines of evidence from independent analyses, we obtained a list of genes that could serve as potential medical targets for mitigating the progression of heart failure in HCM (Figure 2J), and most have not been implicated in heart failure or cardiac hypertrophy such as *FGF12*, *IL31RA*, *BDNF* and *PROS1*. Notably, the expression of *FGF12* (fibroblast growth factor 12) decreased along the trajectory towards the failing state in HCM (Figure 2K). FGF12 has recently been reported to inhibit the pathological remodeling of SMCs in pulmonary arterial hypertension^37^. Likewise, we speculated that it may play a protective role in the pathological remodeling of cardiomyocytes in HCM.

Cardiac fibrosis is a scarring process in the cardiac tissue characterized by excessive deposition of ECM in response to pathophysiological stimuli.^38^ A high burden of cardiac fibrosis exists in HCM patients,^39^ which leads to diastolic dysfunction. Cardiac fibrosis has been proved to be an independent predictor of adverse outcomes including SCD and heart failure in HCM.^40^ Cardiac fibrosis is mediated by the activation of fibroblasts, and understanding the regulatory mechanism underlying the fibroblast activation in HCM is critical for developing effective medical therapies to alleviate the cardiac fibrosis and thereby prevent adverse outcomes for HCM patients. We identified the activated fibroblast subpopulation FB2, which was significantly expanded (Figure 3E) and localized in fibrotic regions (Figure 6E) as expected. Furthermore, based on multiple lines of evidence from independent analyses, we obtained 28 candidate target genes for anti-fibrosis medical development (Figure 3M). Among them, some top-ranked TF genes may represent key regulators driving the fibroblast activation in HCM or other fibrosis-associated conditions. For example, *RUNX1* has recently been suggested to be a key regulator of cardiac fibrosis following myocardial infarction.^10^ A recent study demonstrated that *MEOX1* regulated the pro-fibrotic function and was implicated in the fibrosis of multiple human organs including the heart, liver, lung and kidney.^41^ *AEBP1* (also named *ACLP*) has been implicated in the fibroblast activation of lung fibrosis.^42^ However, our study also identified an array of novel genes and pathways that have not been explicitly implicated in fibrosis. For example, *LEF1* (encoding a transcription factor involved in the Wnt signaling pathway; Figure 3M) and IL18 signaling pathway (Figure 6H). Intriguingly, *NRXN3*, encoding a transmembrane receptor protein of the neurexin family that is predominantly expressed in neurons and is mostly discussed in mental diseases,^43^ was found to be highly expressed in activated fibroblasts (Figure 3B), and its precise role in cardiac fibrosis merit further exploration.

Increasing evidence has shown that immune cells coordinate the responses of cardiomyocytes (e.g., hypertrophy) and other noncardiomyocytes (e.g., fibroblast activation) during pathological cardiac remodeling.^44^ Therefore, identifying the disease-associated immune cell subpopulations and developing therapies regulating the phenotype of cardiac immune cells represent another important strategy for treatment, for example, targeting cardiac fibrosis with engineered T cells.^45^ We explored the alterations of the immune microenvironment in the cardiac tissue of HCM and observed the activation of both innate (e.g., tissue-resident macrophages) and adaptive (e.g., T/NK cells) immunity (Figure 4). Meanwhile, immune response-related pathways, for example, antigen processing and presentation, were found to be upregulated in all the nonimmune cell types, reflecting an enhanced immune response in HCM. The TGF-β signaling has many pleiotropic effects not only in disease, for example, promoting cardiac hypertrophy and fibrosis in the pathological cardiac remodeling, but also in tissue homeostasis.^46^ While TGF-β blockade may be a promising therapeutic strategy, direct and excessive TGF-β inhibition may result in matrix degradation, cardiac dilation and dysfunction.^47^ Though intercellular communication analysis, we found that the top sender of TGF-β changed from MAC2 in HEALTHY to effector T/NK cells in HCM (Figure 5J), which suggest that inhibiting the activation of T/NK cells may attenuate the TGF-β signaling and thereby alleviate the pathological remodeling in HCM while avoiding the deleterious effects of direct TGF-β blockade.

The number of subjects in the healthy group (n=2) was smaller than the HCM group (n=10) in the current study. This may limit the statistical power of comparative analyses. Additional control samples would be included to address this limitation in subsequent studies. In addition, only the TFs, ligands and receptors were considered in the prioritization of candidate targets for subsequent functional studies of our lab; however, other types of molecules, e.g., kinases, may also serve as ideal targets for drug development. We provided the analysis results for all the genes in supplemental tables for further prioritization by the community.

In conclusion, we provided a comprehensive analysis of the lineage-specific regulatory changes in HCM. Our analysis identified a vast array of candidate therapeutic target genes and pathways to prevent or attenuate the pathological remodeling of HCM. Our datasets constitute a valuable resource to examine the cell type-specific expression changes of HCM at single-nucleus and spatial resolution.

## Supporting information

Table III

Table VII

Table VIII

Table XII

Table XIII

## Acknowledgments

We thank Qingzhi Wang, Yang Lu and Yiwei You at the Center of Laboratory Medicine, Fuwai Hospital for technical support in cryosectioning and Masson’s trichrome staining.

## Supplemental Materials

Extended Methods

Data Supplement Figures I-XIII

Data Supplement Tables I-XIII

