## Extended Methods for "Lineage-specific regulatory changes in the pathological cardiac remodeling of hypertrophy cardiomyopathy unraveled by single-nucleus RNA-seq and spatial transcriptomics"

### SUPPLEMENTAL MATERIAL

#### Extended Methods

##### Nucleus isolation

Frozen myocardial tissue was thawed on ice and dissected into small pieces, followed by washing once using cold PBS (Gibco). Tissues pieces were then transferred into a 50-ml Falcon tube containing 30 ml of lysis buffer (0.32 M sucrose, 5 mM CaCl2, 3 mM C4H6MgO4, 0.5 mM EGTA, 10 mM Tris-HCl 8.0, 2 mM EDTA, 1 mM PMSF, 1 mM DTT and 80 U/ml RI). The tissues were homogenized with a T-25 Ultra-Turrax probe homogenized (IKA) at 24,000 rpm for 15 seconds. A glass douncer (40ml) with a tight pestle was used to homogenize the tissue with 10 strokes. After incubation on ice for 10 minutes, the crude nucleus suspension was passed through a 100-μm and 70-μm nylon mesh cell strainer (BD Biosciences), consecutively, followed by spinning down with centrifuge at 700 g for 10 minutes at 4 ℃. Then the supernatant was removed carefully, and the crude nucleus isolate was suspended in 30 ml sucrose buffer (2.1 M Sucrose, 3 mM C4H6MgO4, 10 mM Tris-HCl 8.0, 1 mM PMSF, 1 mM DTT and 80 U/ml RI). Next, the nucleus sample was centrifuged at 13,000 rpm for 60 minutes at 4 ℃ (Beckman Avanti S-25). When the spin was completed, the supernatant was carefully removed. Subsequently, the nucleus pellet was dissolved using 10 ml nucleus suspension buffer (1% BSA, 200 U/ml RI in PBS) and washed once, followed by spinning down with centrifuge at 500 g for 10 minutes at 4 ℃. The nucleus pellet was then suspended in 1 ml nucleus suspension buffer and subjected to further experiments. All procedures were performed below 4 ℃, and RNA inhibitor (RI, 80U/ml) was added to all buffers.

##### Library preparation for snRNA-seq

The prepared single nuclei suspension was loaded on the Chromium Controller (10X Genomics). 3' Gene expression libraries were prepared using Chromium Next GEM Single Cell 3' GEM, Library & Gel Bead Kit v3.1 according to the manufacturer's protocol. The libraries were sequenced on an Illumina NovaSeq 6000 system.

##### Preprocessing and quality control of the snRNA-seq data

To count both the exonic and intronic reads captured by snRNA-seq, a custom "pre-mRNA" reference package was built based on the human reference genome dataset (version: refdata-gex-GRCh38-2020-A) following the 10X Genomics' instructions. The raw sequencing reads were aligned to the "pre-mRNA" reference using the official toolkit Cell Ranger (v4.0.0). The output nucleus-gene expression matrix was imported into Seurat (v3.2.3)[1,2] for further data preprocessing. To exclude genes that were likely detected from random noise, genes with counts in fewer than 3 nuclei were filtered out. To remove poor-quality nuclei that might have resulted from doublets or other technical noise, nuclei were filtered for UMI counts (500 < nCount_RNA < 50,000), genes (300 < nFeature_RNA < 7000), the proportion of mitochondrial genes (percent.mito < 0.05) and the proportion of ribosomal genes (percent.ribo < 0.05). To further remove possible doublets, nuclei with the doublet scores > 0.35 predicted by Scrublet[3] were filtered out. In addition, nuclei that were enriched in the expression of marker genes for multiple lineages were excluded from further analyses.

##### Normalization, feature selection, integration, scaling and clustering of the snRNA-seq data

For each sample, the sum of the UMI counts for each nucleus was normalized to 10,000, and then the normalized UMI counts were log-transformed. For each sample, 2,000 genes were selected using the "FindVariableFeatures" function of Seurat. To correct for potential batch effects and identify shared cell states across samples, we integrated nuclei of all samples via canonical correlation analysis (CCA) implemented in Seurat. To mitigate the effects of unwanted sources of variation, we regressed out the proportion of mitochondrial genes, UMI count, gene number, the proportion of mitochondrial genes and the proportion of ribosomal genes with linear models using the "ScaleData" function. Next, the data were centered for each gene by subtracting the average expression of that gene across all nuclei, and then were scaled by dividing the centered expression by the standard deviation. The scaled data was then subjected to linear dimensional reduction through principal component analysis (PCA). The first 30 PCA components were used to compute a shared nearest neighbor (SNN) graph of the nuclei. The SNN graph was embedded in two-dimensional space using a non-linear dimensional reduction method, i.e., uniform manifold approximation and projection (UMAP).[4] The clustering of all nuclei was performed using the Louvain algorithm.[5]

**Identification of the differentially expressed genes in a specific cell type based on the snRNA-seq data**

To detect the differentially expressed genes in a specific cell type between HCM and HEALTHY, we performed differential expression analysis using a method implemented in the R package DEsingle,[6] which employs a zero-inflated negative binomial model to estimate the fraction of dropout and real zeros in snRNA-seq data. A gene was significantly differentially expressed if it met the following criteria: the absolute of log2 fold change >1, adjusted P-value < 0.05, and being categorized as "general differential expression" that means that the gene had significant differences in both the expression abundances and the fractions of real zeros between HCM and HEALTHY.

**Differential proportion analysis based on the snRNA-seq data**

To determine whether the change in the relative proportion of a specific lineage or cluster in HCM was expected by chance compared with HEALTHY, we performed a permutation-based statistical test (differential proportion analysis; DPA) as described previously.[7] The statistical threshold was set to be a P-value < 0.05.

**Gene set enrichment analysis based on the snRNA-seq data**

All the expressed genes in the snRNA-seq data were pre-ranked by Signal2Noise (the difference of means between HCM and HEALTHY scaled by the standard deviation). Then, the ranked gene list was imported into the GSEA software (version: 4.0.1).[8] A nominal P-value < 0.05 and an FDR q-value < 0.25 were set to be statistically significant. The precompiled canonical pathway gene sets ("c2.cp") in MSigDB (version: 7.2)[9] were used in this analysis.

**Differential gene regulatory network analysis based on the snRNA-seq data**

Gene regulatory networks (GRNs) in a specific lineage were built based on the single-nucleus datasets and comparative analysis of the GRNs between HCM and HEALTHY was performed using the method implemented in bigScale2.[10] Briefly, the GRN of a specific lineage was inferred with the 'compute.network' function separately for each condition. The number of edges of the inferred GRNs was homogenized throughout the networks using the 'homogenize.networks' function. Finally, changes in node centralities (the relative importance of genes in the network) in HCM compared to HEALTHY were identified using the 'compare.centrality' function. Gene rankings based on the changes in centrality were output separately for each of the four measures of centrality including degree, betweenness, closeness and pagerank. The networks were visualized with Cytoscape (version: 3.7.0).

**Trajectory inference based on the snRNA-seq data**

Trajectory inference was performed to order the nuclei along a biological process of interest, e.g., fibroblast activation, using Slingshot (v1.4.0)[11] under default settings. To increase the robustness of the inference, the combined snRNA-seq data of HCM and HEALTHY was considered. A Kolmogorov-Smirnov test was performed to assess whether differences existed between the pseudotime distributions of the two conditions. Then, the genes with different expression patterns along the trajectory between the two conditions were identified using tradeSeq (v1.6.0).[12] For each condition, a smoothed expression profile along the inferred trajectory was estimated for each gene, using a negative binomial generalized additive model (the function “fitGAM”, nknots = 5). Then, the fitted model was used as an input to the “conditionTest” function to test whether genes have different expression patterns along the trajectory between conditions (referred to as differential expression pattern analysis). The significance threshold was set to be a P-value adjusted for multiple testing < 0.05.

**Ligand-receptor interaction analysis based on the snRNA-seq data**

Based on the snRNA-seq data, CellChat (v0.5.5)[13] was used to infer ligand-receptor interactions among subpopulations separately for each condition, and to identify the signaling changes in HCM through comparative analysis by following tutorials of the software. Briefly, overexpressed ligands or receptors for each subpopulation were detected. Next, each potential communication between any two subpopulations was quantified by a communication probability (interaction strength) value, which was modeled by the law of mass action. Significant interactions (P-value < 0.05) were determined by a permutation test that randomly permuted the subpopulation labels and then recalculated the communication probability. By leveraging pattern recognition approaches, dominant incoming and outgoing signal patterns for each subpopulation were detected in HCM or HEALTHY. Major signaling sources and targets of the signaling network for a specific pathway were inferred through network centrality analysis. Joint manifold learning of the HCM and HEALTHY communication networks was performed to group the signaling pathways based on functional similarity (a high degree of functional similarity means that the major senders and receivers are similar). Then, the signaling pathways with large changes in terms of functional similarity in HCM, which means that the major senders and receivers changed for the signaling pathways, were identified based on the Euclidean distance in the learned joint manifold. The conserved or greatly changes signaling pathways in HCM were identified by comparing the overall information flow (the sum of communication probability among all pairs of cell groups in the inferred network) of each signaling pathway in HCM versus HEALTHY.

**Sample preparation for spatial transcriptomic assays**

The RNA quality of the OCT-embedded cardiac IVS tissue block was measured using an Agilent 2100 bioanalyzer. The tissue blocks with an RNA integrity number (RIN) greater than 6 were used for 10X Visium spatial transcriptomic assays. Cryosectioning was performed on a Leica CM3050 S cryostat and brightfield images were taken on a Leica Aperio VERSA whole-slide scanner at 20× resolution.

**Tissue optimization for spatial transcriptomic assays**

The optimization of the conditions for tissue permeabilization was performed according to 10X Visium Spatial Tissue Optimization User Guide (CG000238, 10X Genomics). Briefly, tissue sections were placed on the capture areas of a tissue optimization slide. The sections were fixed, stained, and then permeabilized for different times. The mRNA released during permeabilization bound to oligonucleotides on the capture areas. Fluorescent cDNA was then synthesized on the slide and was imaged. The optimal permeabilization time was the one that resulted in the maximum fluorescence signal and the lowest signal diffusion.

**Sequencing libraries preparation for spatial transcriptomic assays**

Sequencing libraries were constructed using the Visium Spatial Gene Expression Slide & Reagent kit (10X Genomics) following the manufacturer’s instructions. Briefly, a 10-μm frozen tissue section was placed on one of the capture areas (6.5 × 6.5 mm with ~5000 barcoded spots) of a gene expression slide. Following Hematoxylin and Eosin (H&E) staining, brightfield images were taken. Tissue permeabilization was performed for the optimal time, as determined in the tissue optimization procedure. Then, reverse transcription was done and sequencing libraries were prepared. Sequencing was conducted using an Illumina NovaSeq 6000 system.

**Processing of the spatial transcriptomic data**

Sequencing read alignment, fiducial/tissue detection, and spot barcode/UMI counting of the spatial transcriptomic data were performed separately for each section using the 10X Genomics official tool kit Space Ranger (v1.2.2), which takes a H&E-stained brightfield image and fastq files as inputs. The same version of the human reference genome dataset as that used in the processing of snRNA-seq data was applied, i.e., refdata-gex-GRCh38-2020-A. The output gene-spot matrix was imported to Seurat for downstream analysis and visualization. Only spots that have been determined to be over tissue were retained. To account for technical artifacts while preserving biological variance in UMI counts across sports, the normalization method sctransform[14] was used. Linear dimensional reduction with PCA was performed using the function "RunPCA". An SNN graph was constructed based on the first 30 PCA components using the function "FindNeighbors". Based on the SNN graph, the Louvain algorithm was used to cluster the spots (resolution: 0.4) using the function "FindClusters". Finally, UMAP dimensional reduction was performed using the function "RunUMAP" for visualizing the spots in a 2D space.

**Identification of the molecular signature for each nucleus cluster/spot cluster**

The molecular signature of each nucleus/spot cluster was obtained by comparing the transcriptome of each cluster with that of the other clusters using a likelihood-ratio test (test.use: ''bimod'') implemented in the ''FindMarkers'' function of Seurat. The significance threshold was set to an adjusted P-value < 0.05 and a log2 fold change > 0.25.

**Pathway activity scoring in each nucleus/spatial transcriptomic spot**

To quantify the expression activity of a given gene set/pathway in each nucleus/ST spot, a gene set/pathway activity score was calculated for each nucleus or spot using the method implemented in Single Cell Signature Explorer.[15] The precompiled canonical pathway gene sets ("c2.cp") in MSigDB were used in this analysis. The expression activity of a pathway in each sample was represented by the mean of the computed scores across the nuclei/spots in the sample. Based on the calculated pathway activity score of each spatial transcriptomic spot, the differentially regulated pathways of the spots in fibrotic versus non-fibrotic regions of cardiac tissue sections were detected using the Wilcoxon rank-sum test implemented in the function "FindMarkers" of Seurat.

**Integration of the spatial transcriptomic data with the snRNA-seq data**

To integrate with the snRNA-seq data and predict the underlying cellular composition for each spot that contained multiple nuclei, we applied the label transfer workflow of Seurat to assign each spot prediction scores for the subpopulations obtained from the snRNA-seq data analysis.

**Functional enrichment analysis**

Functional enrichment analyses of a list of genes were performed using ClueGO[16] with a Bonferroni corrected P-value threshold of 0.05. Databases including Gene ontology biological process, REACTOME and KEGG were considered in this analysis.

**Masson's trichrome staining**

To histologically assess cardiac fibrosis, Masson's trichrome staining was conducted on the tissue sections adjacent to those used for spatial transcriptomic assays. Masson's Trichrome stain kit (Beijing Yili Fine Chemicals Co., Ltd.) was used by following the manufacturer's protocol. In brief, the frozen sections were incubated in 0.5% hydrochloric acid alcohol for 5 seconds, and then were stained in Weigert's Iron Hematoxylin Solution for 2 minutes. Next, they were stained in Ponceau S fuchsin for 5 minutes. Subsequently, they were incubated in phosphomolybdic acid solution for 2 minutes and were stained in Brilliant Green solution for 2 seconds. Finally, they were incubated in 1 % Glacial acetic acid for 1 minute and then were placed in xylene for 5 minutes. Images were captured using a Pannoramic SCAN II scanner (3DHISTECH).


#### Supplemental table legends

**Table I. Demographic and clinical information of subjects recruited in this study.**

**Table II. Quality metrics of the snRNA-seq datasets.**

**Table III. Quality metrics of the spatial transcriptome datasets.**

**Table IV. The mean normalized expression of all genes in each cell type under HCM or HEALTHY conditions.**

**Table V. Gene signature of each subpopulation in each cell type (multiple sheets).**

**Table VI. Differentially expressed genes of each cell type in HCM versus HEALTHY (multiple sheets).**

**Table VII. Functional enrichment analysis of the upregulated genes of each cell type in HCM versus HEALTHY (multiple sheets). Neuronal cells, lymphatic endothelial cells and T/NK cells are not considered in this analysis due to the limited number of cells.**

**Table VIII. Differentially regulated pathways in HCM versus HEALTHY detected by gene set enrichment analysis (multiple sheets). Neuronal cells, lymphatic endothelial cells and T/NK cells are not considered in this analysis due to the limited number of cells.**

**Table IX. Differential regulatory network analysis of each cell type in HCM versus HEALTHY (multiple sheets).**

**Table X. Genes with differential expression patterns along the pseudotime between HCM and HEALTHY in cardiomyocytes (sheet1) and fibroblasts (sheet2) detected by tradeSeq.**

**Table XI. Ligand-receptor interactions in cardiac tissue of HCM (sheet1) and HEALTHY (sheet2) inferred by CellChat.**

**Table XII. Differentially expressed genes of 10X Visium spatial transcriptome spots in fibrotic versus non-fibrotic regions of cardiac tissue sections from HCM patients (multiple sheets).**

**Table XIII. Differentially regulated pathways of 10X Visium spatial transcriptome spots in fibrotic versus non-fibrotic regions of cardiac tissue sections from HCM patients (multiple sheets).**

#### Supplemental Figures and Figure Legends

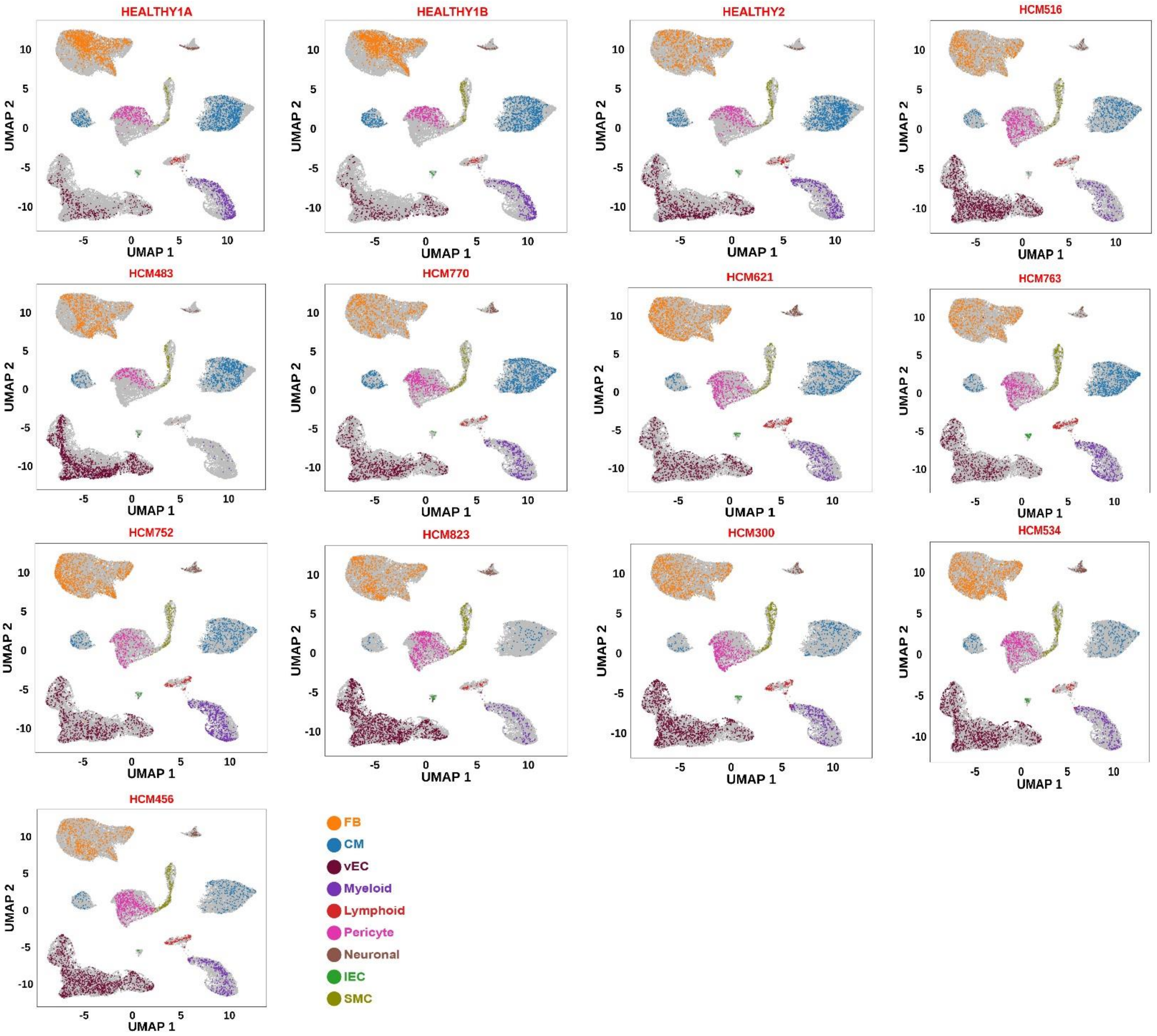


**Figure I. Distribution of each cell type in the UMAP space for each sample. For comparison, an equal number of cells (3,169) are shown for each sample.**

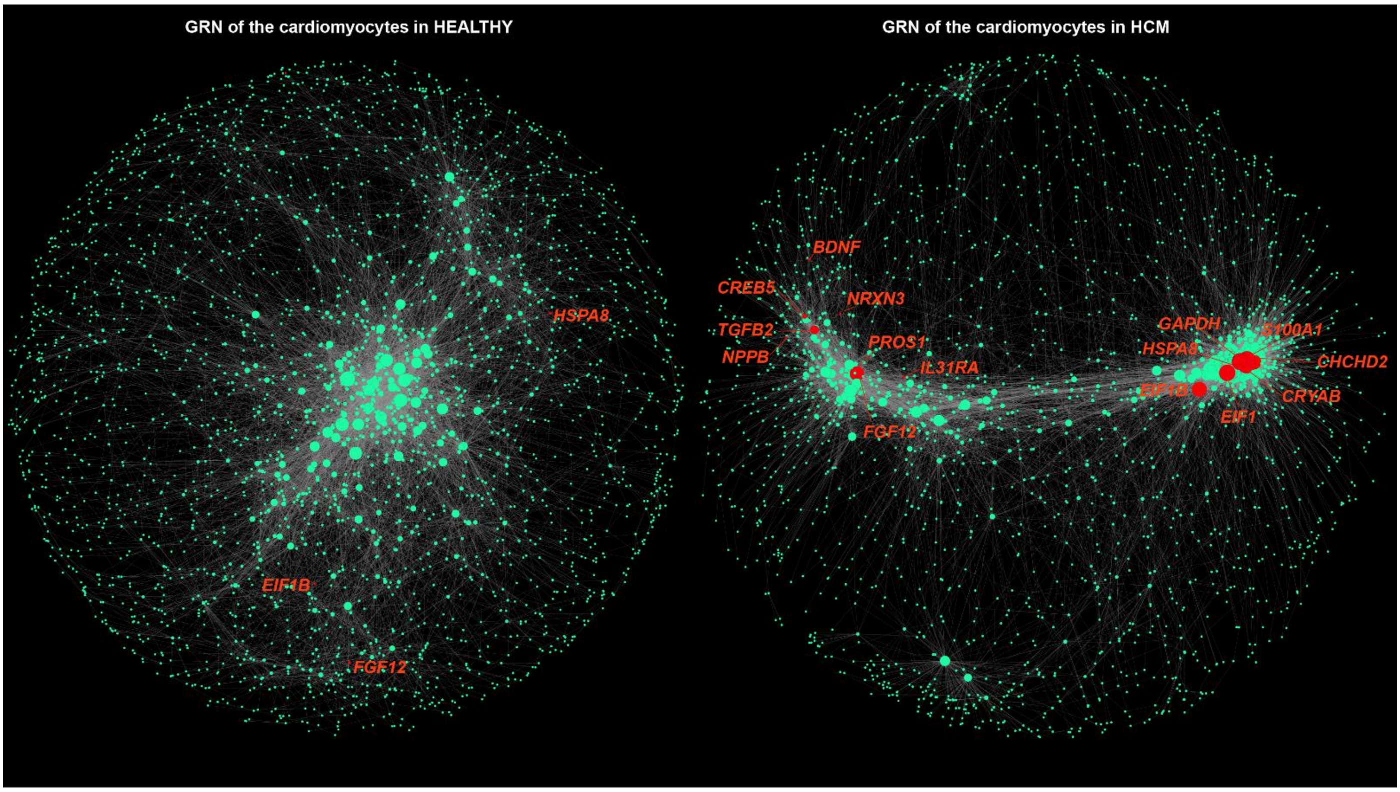


**Figure II. Comparative analysis of the GRNs of the cardiomyocytes between HEALTHY (left panel) and HCM (right panel).** The node size reflects degree centrality. Nodes in red are representative genes with increased biological importance in the network of HCM compared with that of HEALTHY.

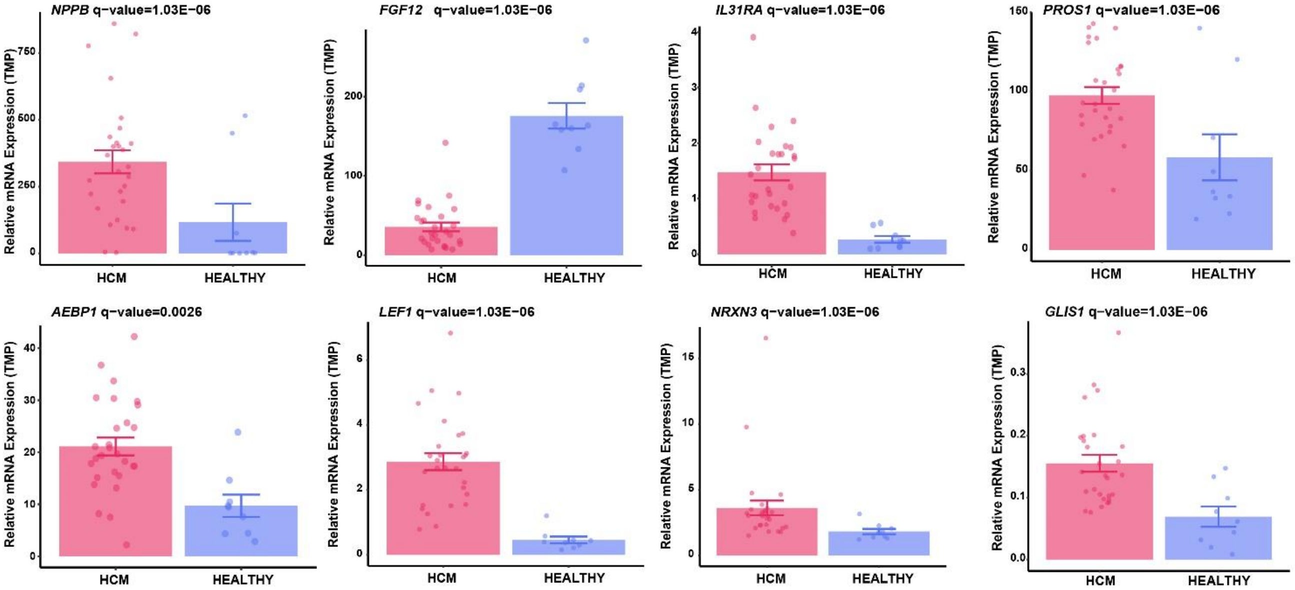


**Figure III. Relative mRNA expression of representative candidate target/marker genes in HCM (n=28) and HEALTHY (n=9) determined by bulk RNA-seq.**

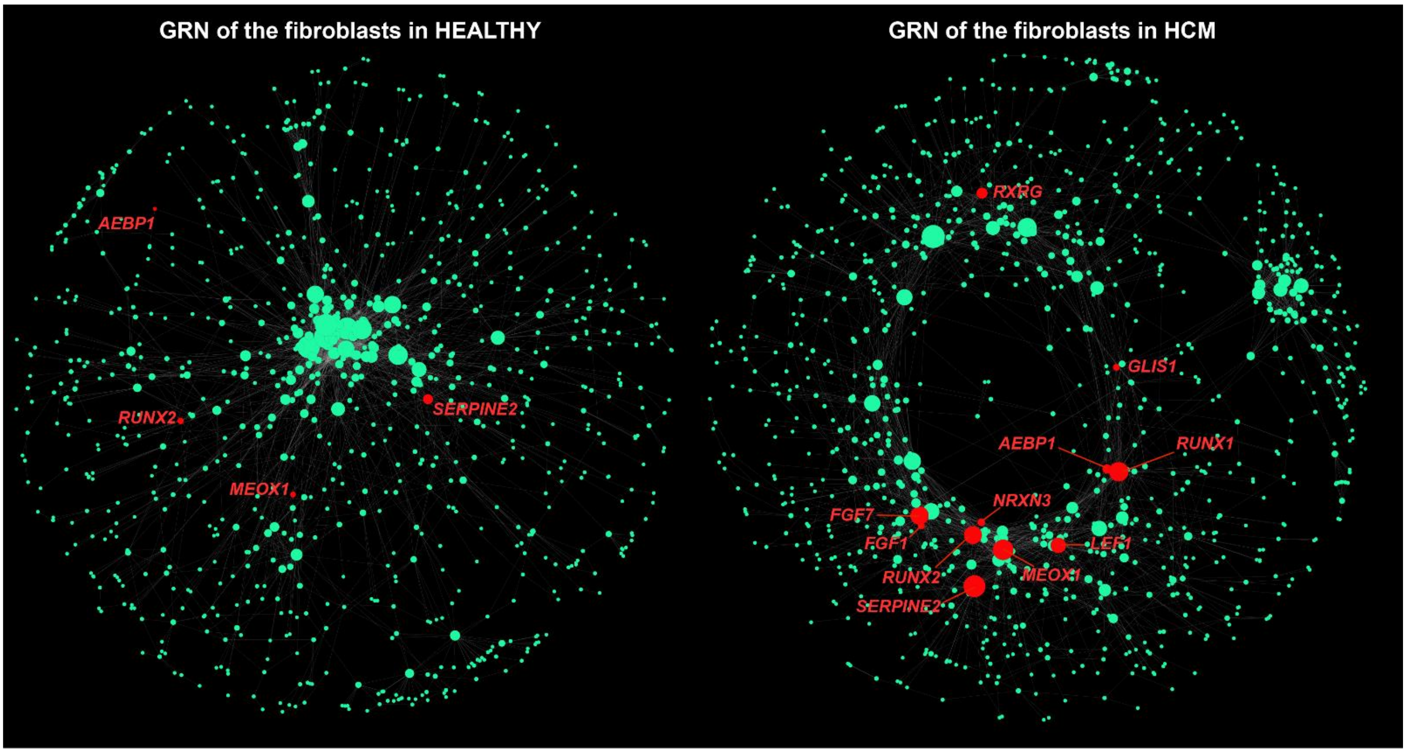


**Figure IV. Comparative analysis of the GRNs of the fibroblasts between HEALTHY (left panel) and HCM (right panel).** The node size reflects degree centrality. Nodes in red are representative genes with increased biological importance in the network of HCM compared with that of HEALTHY.

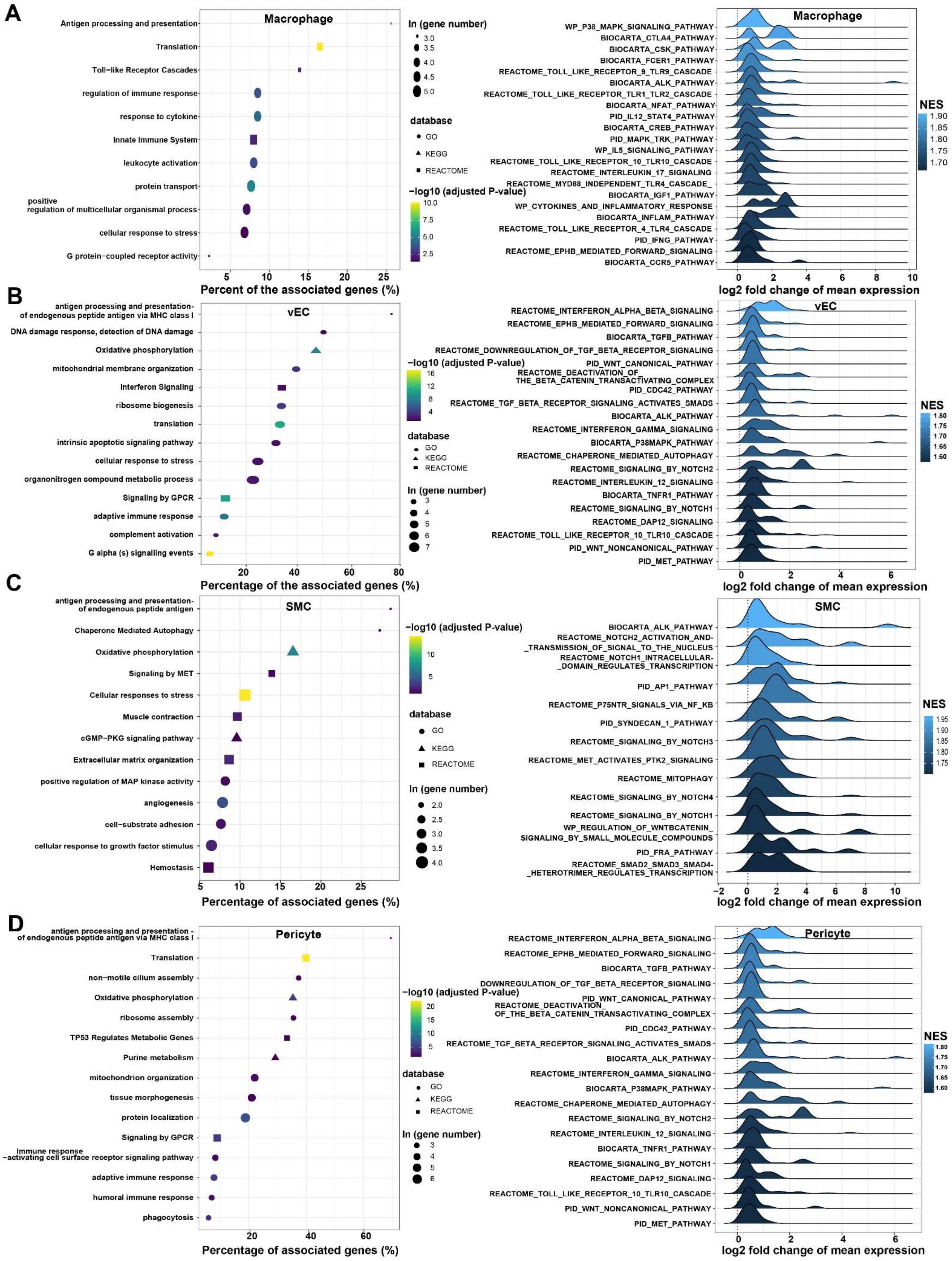

**Figure V. Functional enrichment of the upregulated genes (left panel) and the upregulated pathways revealed by GSEA in macrophages (A), vascular endothelial cells (B), smooth muscle cells (C) and pericytes (D).**

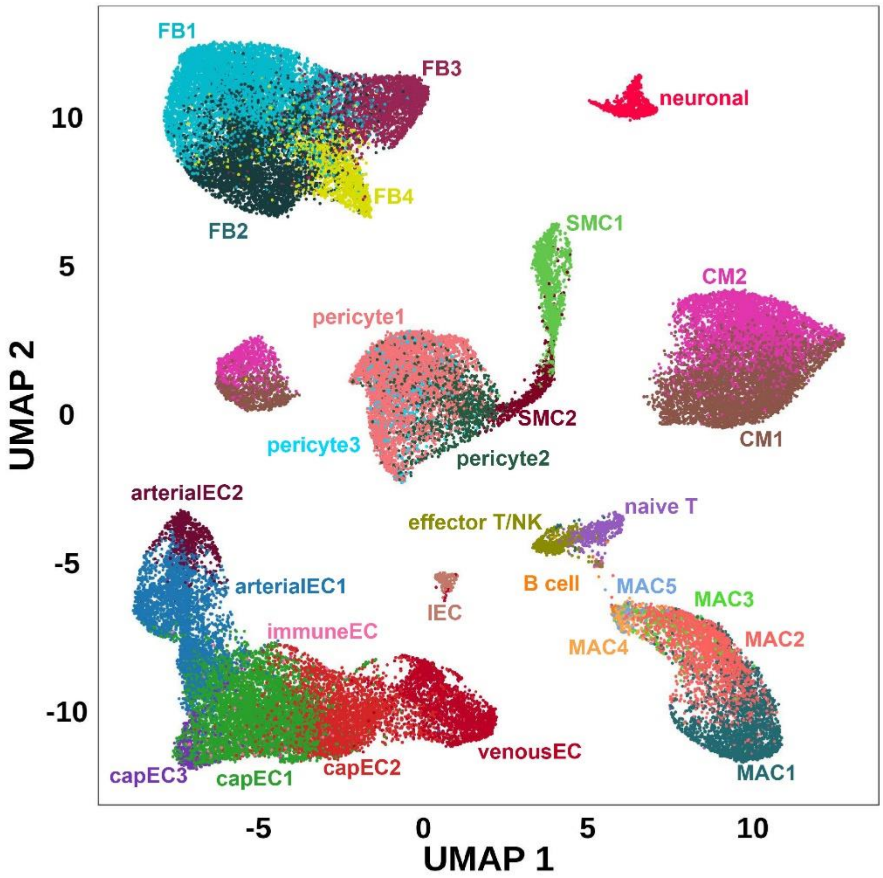


**Figure VI. The distribution of each subpopulation in the UMAP space.**

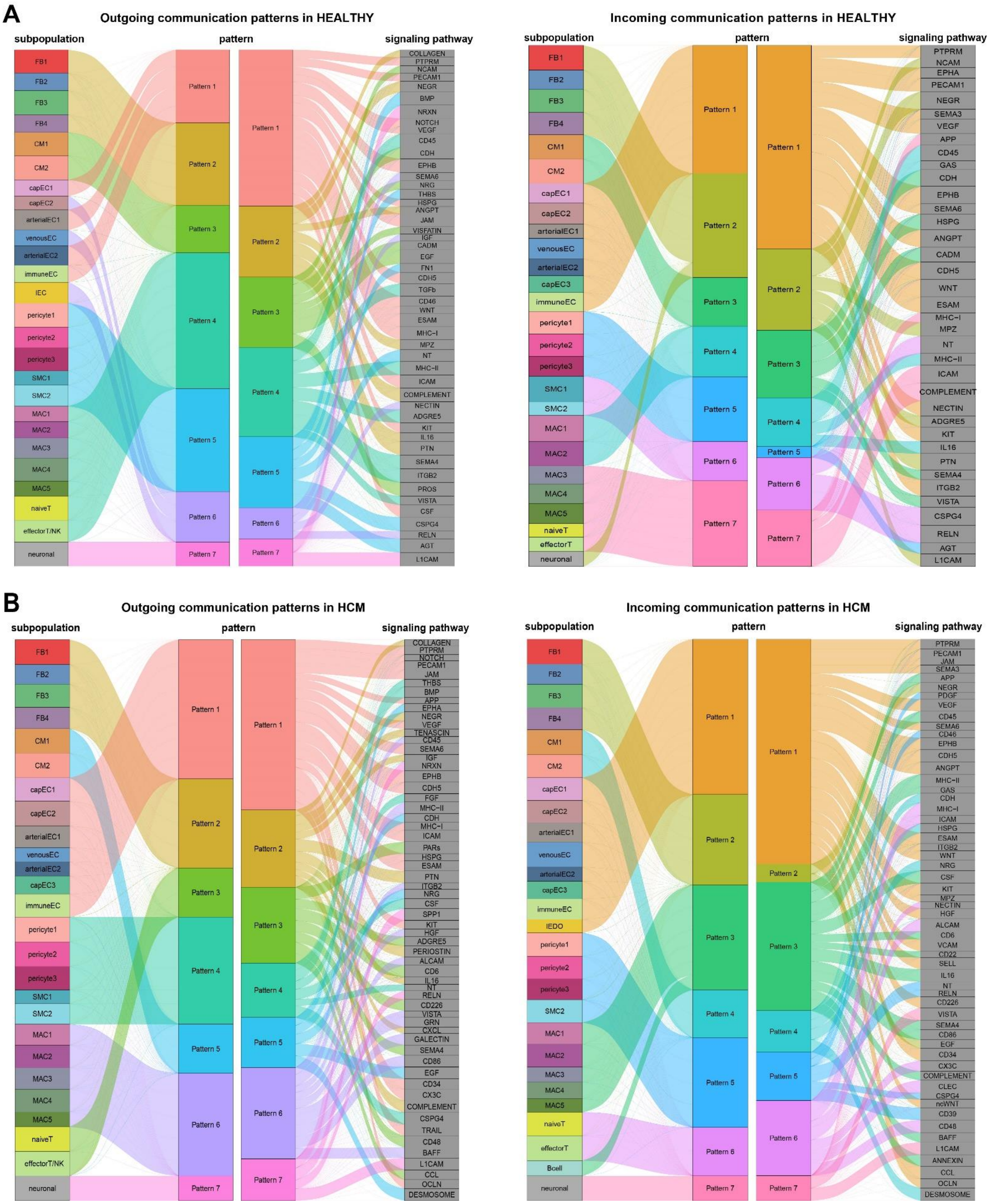


**Figure VII. River plot showing the patterns of the inferred outgoing (left panel) and incoming (right panel) signaling pathways for each subpopulation in HEALTHY (A) and HCM (B).** The thickness of the flow indicates the contribution of the subpopulation or signaling pathway to each pattern.

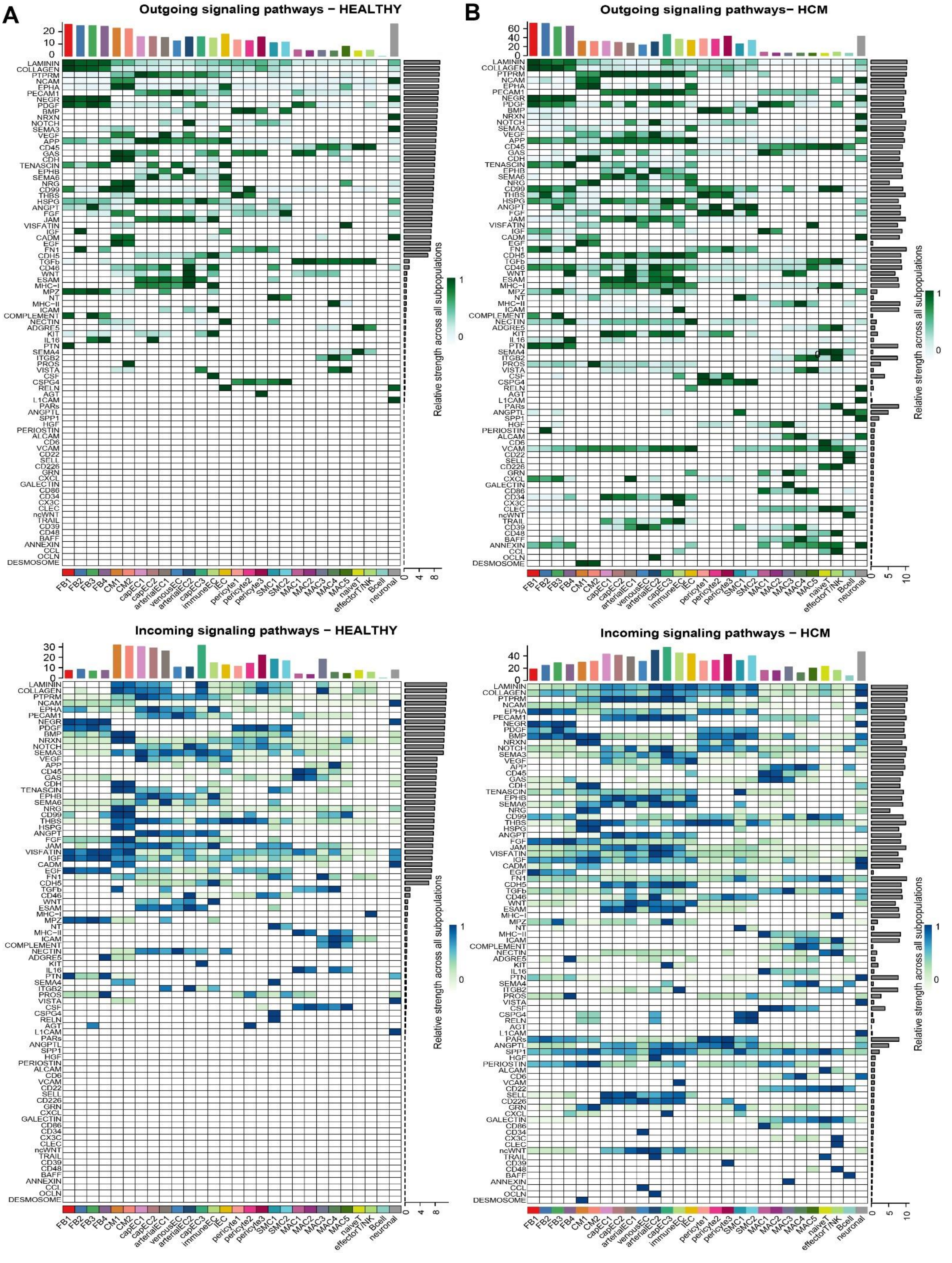


**Figure VIII. Heatmap showing the relative signaling strength of the inferred outgoing (upper panel) and incoming (lower panel) signaling pathways for each subpopulation in HEALTHY (A) and HCM (B).** The top bar represents the sum of the relative signaling strength of all pathways in each subpopulation. The right bar represents the aggregated signaling strength of a signaling pathway across all subpopulations.

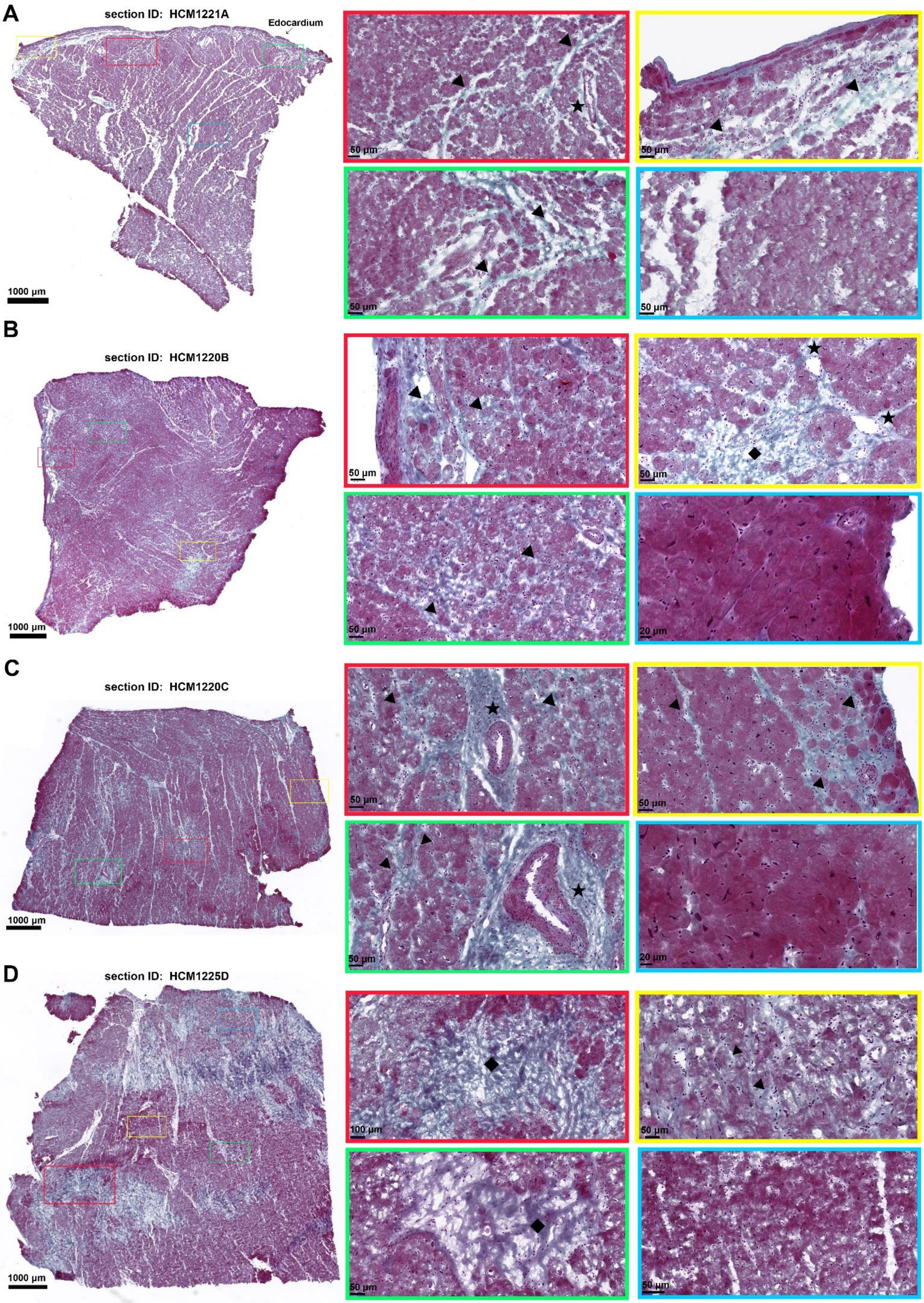


**Figure IX. Masson's trichrome staining showing fibrosis on the cardiac sections adjacent to the four**

**sections selected for spatial transcriptomic assays. (A)** Masson's trichrome staining for the section adjacent to HCM1221A. **(B)** Masson's trichrome staining for the section adjacent to HCM1220B. **(C)** Masson's trichrome staining for the section adjacent to HCM1220C. **(D)** Masson's trichrome staining for the section adjacent to HCM1225D. Fibrotic regions are indicated by collagens stained in blue. ◆: replacement fibrotic scar; ★ perivascular fibrosis; ▲: diffuse interstitial fibrosis including fibrous bands surrounding the cardiac muscle bundles or individual cardiomyocytes. The small panels with cyan border show cells in non-fibrotic regions.

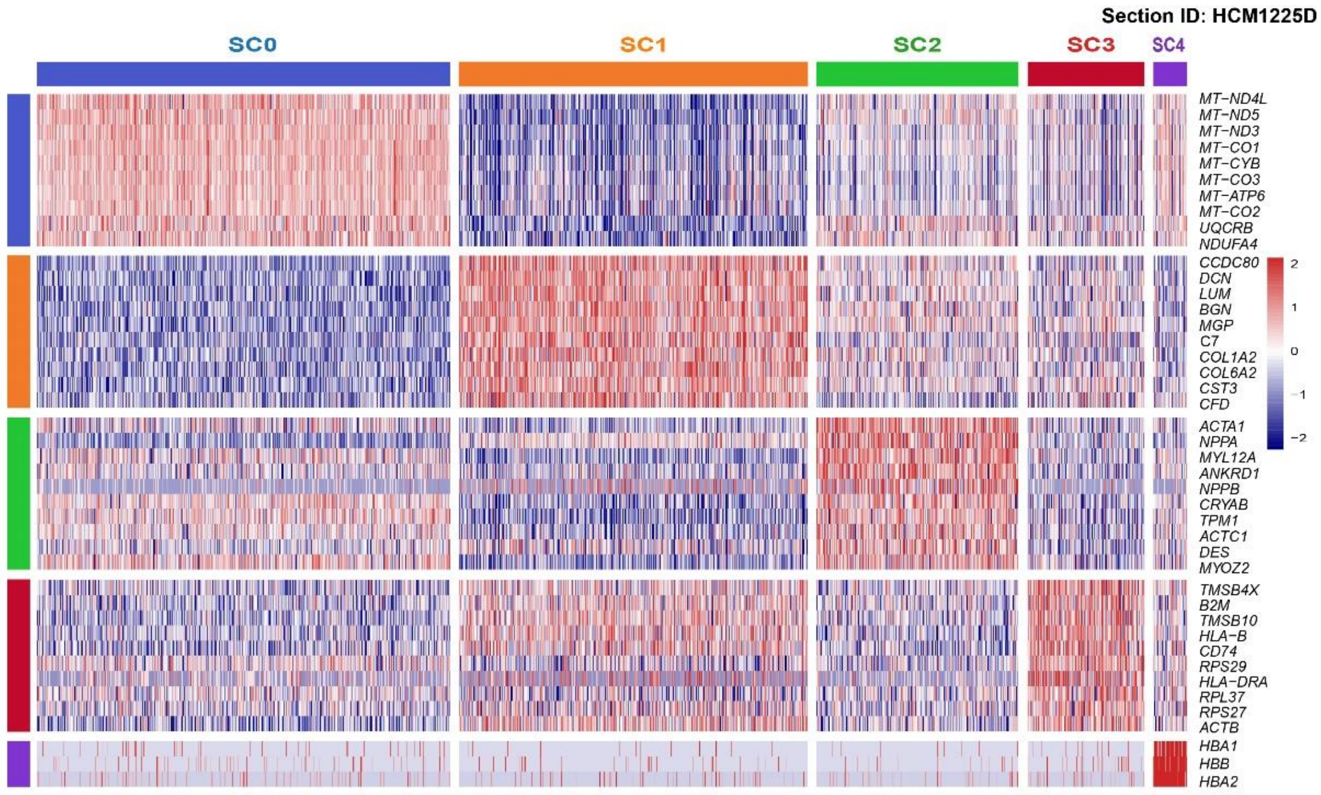


**Figure X. Heatmap showing the expression signature of each spot cluster for the cardiac tissue section HCM1225D.**

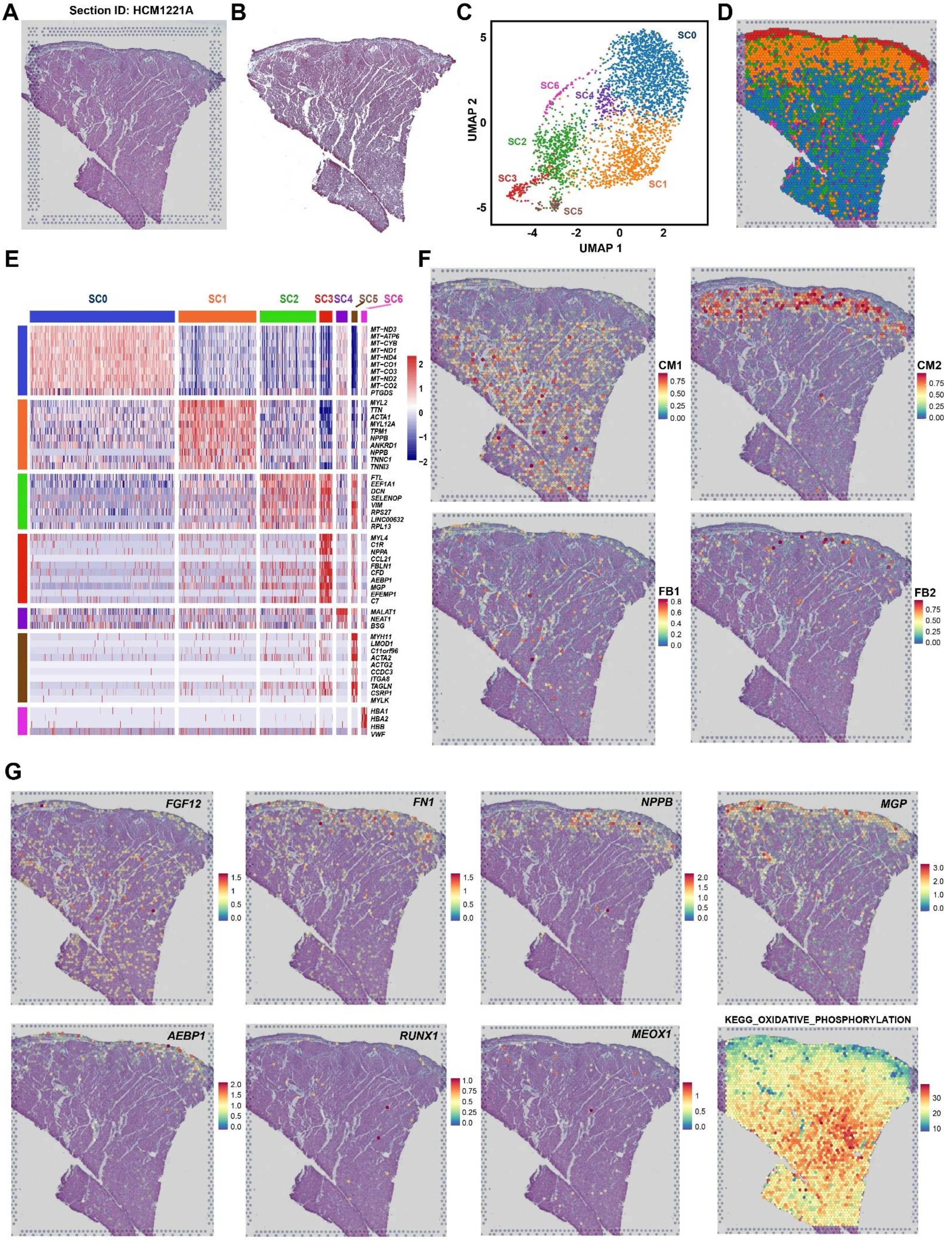


**Figure XI. Spatial transcriptomic analysis of the cardiac tissue section HCM1221A. (A)** H&E staining image. **(B)** Masson's trichrome staining image. **(C)** Clustering of the ST spots. **(D)** Spots color-coded by spot clusters. **(E)** Heatmap showing the expression signature of each spot cluster. **(F)** Probabilistic classification of each spot for the subpopulation CM1, CM2, FB1 and FB2. The integration of snRNA-seq and spatial transcriptomic data was performed by following the label transfer workflow of Seurat. **(G)** The expression distribution of representative markers, candidate genes and pathways on the section.

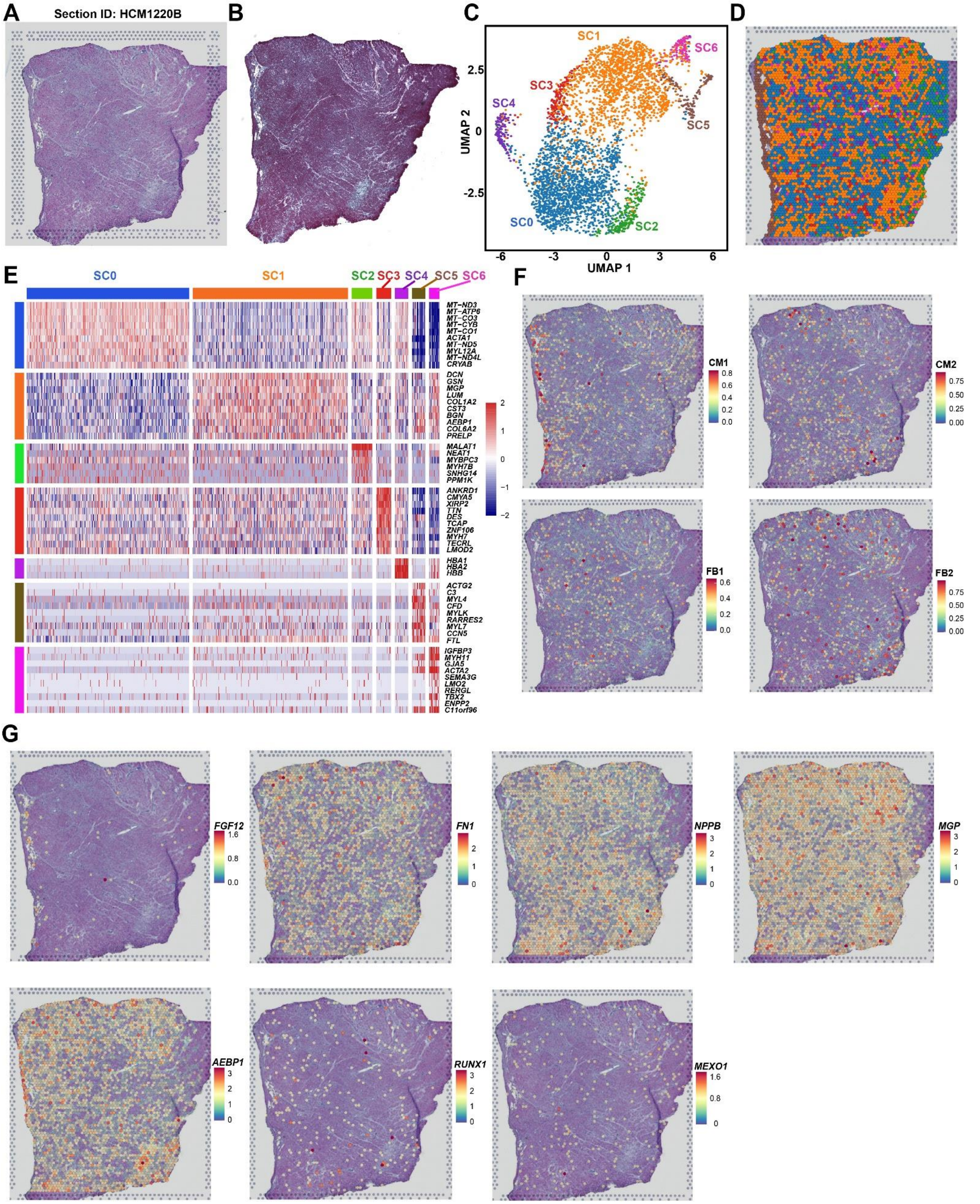


**Figure XII. Spatial transcriptomic analysis of the cardiac tissue section HCM1220B. (A)** H&E staining image.

**(B)** Masson's trichrome staining image. **(C)** Clustering of the spots. **(D)** Spots color-coded by spot clusters. **(E)** Heatmap showing the expression signature of each spot cluster. **(F)** Probabilistic classification of each spot for the subpopulation CM1, CM2, FB1 and FB2. The integration of snRNA-seq and ST data was performed by following the label transfer workflow of Seurat. **(G)** Expression distribution of representative markers and candidate genes on the section.

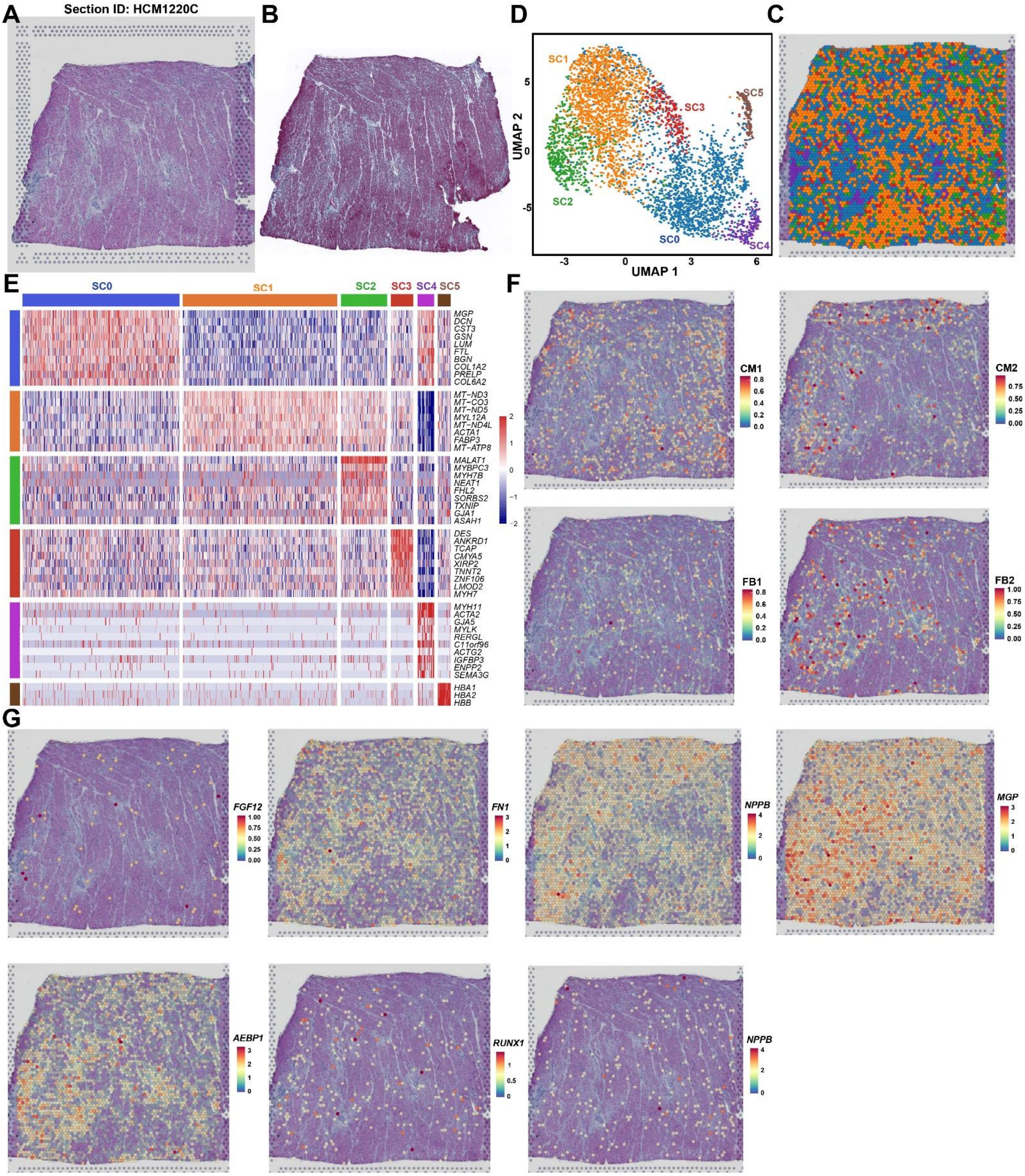


**Figure XIII. Spatial transcriptomic analysis of the cardiac tissue section HCM1220C. (A)** H&E staining image. **(B)** Masson's trichrome staining image. **(C)** Clustering of the spots. **(D)** Spots color-coded by spot clusters. **(E)**

Heatmap showing the expression signature of each spot cluster. **(F)** Probabilistic classification of each spot for the subpopulation CM1, CM2, FB1 and FB2. The integration of snRNA-seq and ST data was performed by following the label transfer workflow of Seurat. **(G)** Expression distribution of representative markers and candidate genes on the section.
